# A Modular Chromosomally Integrated Yeast Surface Display Platform for Quantitative Serology and Immune Profiling

**DOI:** 10.64898/2026.09.08.750026

**Authors:** Dan Lou, Hans Hombauer, Florian Huber, Anna Denzler, Lars Maurer, Konrad Herbst, Matthias Meurer, Hong-Yu Lee, Daniel Kirrmaier, Ivonne Morales, Edward W. Green, Paul Schnitzler, Uta Merle, Claudia M. Denkinger, Tobias Boettler, Viet Loan Dao Thi, Michael Knop

## Abstract

Precise quantification of antigen-specific antibodies is essential for assessing immune status and fundamental to both clinical and research settings. However, current serological platforms remain limited in their ability to combine quantitative performance, flexibility, and scalability. Here, we present a chromosomally integrated yeast surface display (YSD) system engineered for stable, quantitative, and scalable antibody profiling. Genomic integration of the YSD cassette supports uniform antigen presentation without continuous selective pressure, reducing variability associated with plasmid-based systems. Two distinct cell-wall anchoring strategies are implemented within a modular synthetic framework that enables rapid, cloning-free strain construction. Coupled with flow-cytometric readout, the system provides sensitive detection of antigen-specific antibodies with single-cell normalization for antigen expression. In addition, the combination of barcoded YSD strain libraries with fluorescence-activated cell sorting and next-generation sequencing enables multiplexed serological profiling across multiple samples and antigens. As a proof of concept, we demonstrate quantitative antibody detection against SARS-CoV-2 and extend the approach to hepatitis virus antigens, highlighting the utility of this platform as a flexible framework for serological assay development and immune profiling.

## Introduction

Antibodies are central mediators of adaptive immunity and provide clinically and biologically informative readouts of infection history, vaccination, autoimmune reactivity, therapeutic exposure, and immune protection (Schroeder & Cavacini, 2010; Lu et al., 2018). Precise measurement of antigen-specific antibodies in human samples is therefore essential for infectious disease diagnostics, vaccine evaluation, seroepidemiology, immunomonitoring, autoimmune disease stratification, and the development and monitoring of antibody-based therapeutics (Crescioli et al., 2026; Khoury et al., 2021; Long et al., 2020; Metcalf and Graham, 2018; Paul et al., 2024; Plotkin, 2010). As the need to profile humoral immune responses continues to grow—driven by emerging infectious threats, population-scale serosurveillance, antigenic variation, precision immunotherapy, and the expanding clinical use of antibody-based drugs—there is a clear demand for serological platforms that combine quantitative performance, antigen flexibility, scalability, cost-effectiveness, and straightforward assay standardization.

Conventional antibody detection assays, including enzyme-linked immunosorbent assays (ELISA-Assays; Engvall & Perlmann, 1971), virus neutralization assays, and bead-based immunoassays, are widely used, but each has inherent limitations (Krammer & Simon, 2020). ELISAs have long served as a standard format for serological testing because they are robust, accessible, and scalable, but they typically require purified recombinant antigens immobilized on solid surfaces, which may alter antigen presentation or fail to preserve conformational and multivalent epitopes in their native context (Moritz et al., 2022). In addition, ELISAs are limited in multiplexing capacity and can be time-consuming due to repeated incubation and washing steps. For antiviral antibodies, neutralization assays provide biologically relevant information by measuring the capacity of antibodies to inhibit infection. However, live-virus neutralization assays are labor-intensive, low-throughput, difficult to standardize across laboratories, and often require specialized biosafety infrastructure (Roehrig et al., 2008; Bewley et al., 2021; Muruato et al., 2020). Other approaches, most prominently fluorescence-coded bead-based technologies such as Luminex, enable multiplexed antibody profiling from small sample volumes, but they require specialized instrumentation, recombinant antigens for bead coupling, and assay-specific optimization and validation to control for antigen-dependent coupling efficiency, background binding, cross-reactivity, and multiplex interference (Tait et al., 2009; Stern et al., 2023; Roy et al., 2023). Together, these limitations highlight the need for serological platforms that preserve antigen flexibility while supporting quantitative, scalable, and standardized antibody detection.

Yeast surface display (YSD) technology provides a powerful and versatile platform that addresses many of these constraints (Boder & Wittrup, 1997; McMahon et al., 2018; Starr et al., 2020; Wang et al., 2022). In YSD, antigens are expressed as fusions to cell wall-anchoring proteins on the surface of *Saccharomyces cerevisiae*, enabling eukaryotic folding and secretory pathway-associated post-translational processing that can support near-native conformations (Boder & Wittrup, 1997; McMahon et al., 2018). Surface display of antigens followed by antibody staining and flow cytometry allows quantitative, single-cell measurement of antigen expression and antibody binding with high dynamic range and reproducibility (McMahon et al., 2018; Starr et al., 2020; Wang et al., 2022). However, adapting YSD for serological analysis faces several challenges, including reproducible antigen display, efficient strain construction, and robust normalization across experiments. Most existing YSD platforms rely on episomal plasmid-based expression, which can be affected by plasmid instability, variable copy number, dependence on continuous selective pressure, and labor-intensive cloning workflows (Boder & Wittrup, 1997; McMahon et al., 2018; Starr et al., 2020). These features can constrain scalability and throughput, introduce batch-to-batch variability, and complicate assay standardization, thereby limiting broader implementation in diagnostic and large-scale serological applications.

To overcome these challenges, we developed one-step, chromosomally integrated YSD strategies for two widely used display architectures: the Aga2 system and the glycosylphosphatidylinositol (GPI)-anchor system (Boder & Wittrup, 1997; McMahon et al., 2018). Both were optimized for sensitive and scalable antibody detection in human samples. To this end, we engineered YSD acceptor strains with a modular genetic architecture that enables rapid strain construction through synthetic DNA fragment assembly. This flexible design facilitates the parallel generation of antigen-displaying YSD strains with stable surface expression. When combined with flow cytometry, our approach enables sensitive measurement of antigen-specific antibody binding while simultaneously normalizing for antigen expression at single-cell resolution. We demonstrate the broad applicability of this assay by detecting antibodies against SARS-CoV-2, hepatitis E virus (HEV), and hepatitis C virus (HCV) in human samples. Furthermore, we introduce a multiplexed YSD-based methodology that integrates barcoded YSD strains with fluorescence-activated cell sorting (FACS) and next-generation sequencing to enable parallel profiling of antibody responses across large sample sets. Together, our optimized YSD platform provides a rapid, flexible, and scalable tool for serological analysis and immune profiling.

## Results

### Chromosomally integrated yeast surface display system enables stable and uniform antigen expression

Current YSD platforms are commonly plasmid-based; however, yeast plasmids can be unstable, and even populations maintained under selection can contain substantial fractions of cells that have lost the plasmid, particularly in the case of high-copy-number plasmids (Futcher & Cox, 1984). To establish a robust platform for rapid and uniform YSD, we developed chromosomally integrated YSD systems based on two well-established plasmid-based YSD architectures. The AGA2 platform was adapted from the pYD1 architecture (Boder & Wittrup, 1997), in which the protein of interest (POI) is displayed as an Aga2p fusion protein anchored to the cell wall through disulfide linkage with Aga1p (Figure 1a(i)). The second system was constructed based on the pYDS649 platform (McMahon et al., 2018), in which the POI is tethered to the cell wall via a 649-amino-acid glycosylated stalk domain that is anchored in the plasma membrane through a GPI anchor (Figure 1a(ii)).

**Figure 1.**
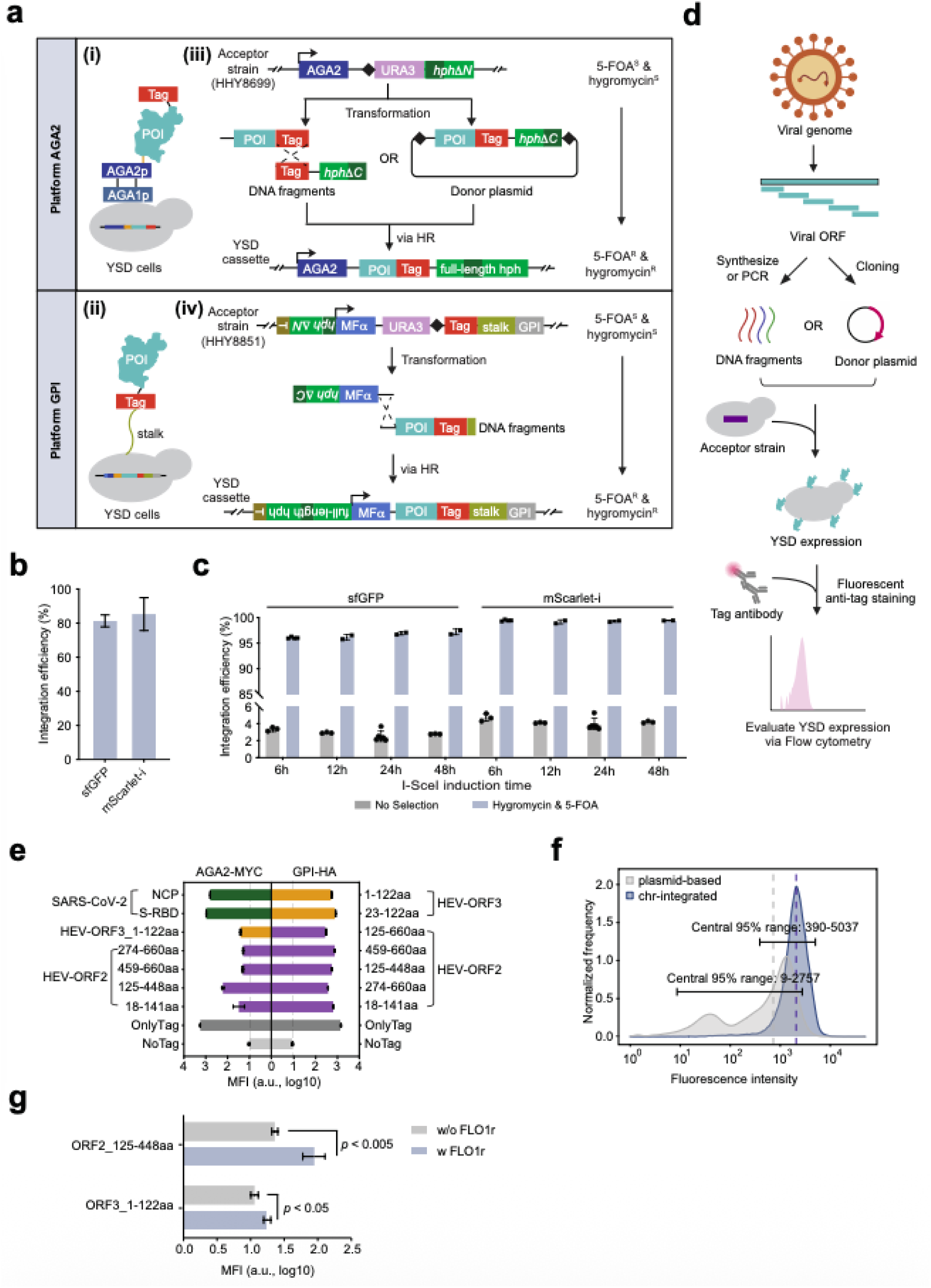
A chromosomally integrated yeast surface display (YSD) system enables stable and uniform antigen expression. (a) i, Illustration of antigen display on the yeast cell surface using the AGA2 platform. ii, Illustration of antigen display on the yeast cell surface using the GPI platform. iii, Schematic of the chromosomal integration strategy for the AGA2 platform. iv, Schematic of the chromosomal integration strategy for the GPI platform. The I-SceI motif is indicated by a black diamond. (b) Integration efficiency of the fragment-based chromosomal YSD construction strategy, assessed using fluorescent protein reporters. Efficiency is shown as the percentage of fluorescent colonies following 5-FOA and hygromycin selection. (c) Validation of the donor plasmid transformation method using sfGFP and mScarlet-i fluorescent reporters. Integration efficiency, defined as the percentage of fluorescent cells after 5-FOA and hygromycin selection, was analyzed by flow cytometry. (d) Overview of the experimental workflow for generating chromosomally integrated YSD strains. (e) Surface expression levels of various antigens displayed using the AGA2-based and GPI-based platforms. Antigens were derived from SARS-CoV-2 and hepatitis E virus (HEV). Strains expressing only the epitope tag served as positive controls, and strains lacking any tag served as negative controls. Median fluorescence intensity (MFI; log10-transformed) of the epitope tag is shown. (f) Surface expression profiles of plasmid-based and chromosomally integrated YSD constructs measured by flow cytometry. Dashed vertical lines indicated the median fluorescence intensity for each group. Horizontal lines indicate the 2.5th - 97.5th percentile range, representing the central 95% of observed fluorescence values. (g) Influence of the FLO1r domain on surface expression of GPI-HEV constructs. Median fluorescence intensity (MFI; log10-transformed) of the epitope tag is shown.

We designed platform-specific generic acceptor strains to enable efficient genomic integration and modular construct assembly (Figure S1). Each acceptor strain harbors a GAL1pr-I-SceI construct, enabling galactose-inducible expression of the I-SceI endonuclease and thereby enhancing integration efficiency by homologous recombination (Plessis et al., 1992). The acceptor module, integrated at the *HOM3* locus, contains a partial YSD cassette, an I-SceI recognition site, a *URA3* cassette, and a truncated hygromycin resistance gene (hph) (Figure 1a; Meurer et al., 2018). In the AGA2 platform, the strains additionally carry an *AGA1* overexpression construct, because Aga1 serves as the cell-wall anchor for the Aga2-POI fusion via disulfide bridges. Integration of the acceptor module disrupts *HOM3*, rendering the cells auxotrophic for threonine and methionine. As this marker is rarely used in subsequent strain construction, the resulting auxotrophy provides a convenient phenotypic readout for rapid validation of correct acceptor-strain generation.

This modular design supports two assembly strategies. In the first strategy, two linear DNA fragments are transformed into yeast cells containing the integrated acceptor module after induction of the I-SceI endonuclease. One fragment encodes the POI, and the other is a generic fragment containing one part of the selection marker. The two fragments integrate into the acceptor locus through overlapping homologous sequences. Correct integration replaces the counter-selectable *URA3* marker originally present at the acceptor locus and reconstitutes a functional *hphNT2* marker from the two partial marker fragments (*hphΔN* and *hphΔC*), which are contained in the acceptor module and the introduced fragment, respectively. This enables efficient selection for correct integration events using positive and negative selection simultaneously (Figure 1a(i, iii)). The second strategy employs a plasmid-borne POI that is first transformed into yeast, followed by I-SceI induction which then triggers genomic integration, followed by selection for correctly integrated clones. For the GPI platform we only implemented the fragment-based strategy (Figure 1a(ii)).

We tested the efficiency of both assembly strategies for the AGA2 platform using fluorescent proteins as model POIs. In the AGA2 platform, the fragment-based approach yielded >80-85% sfGFP-positive and mScarlet-i-positive fluorescent colonies, indicating efficient recombination (Figure 1b; Pédelacq et al., 2006; Bindels et al., 2017). Transformation with donor plasmids carrying either sfGFP or mScarlet-i achieved > 95% correct integrations regardless of I-SceI induction time (Figure 1c).

Notably, the fragment-based approach is readily scalable using synthetic DNA fragments and PCR, thereby minimizing cloning requirements and enabling high-throughput, parallel generation of YSD constructs. In contrast, the plasmid-borne approach can be used for screening large variant libraries, for example when mutagenized protein libraries are generated directly in yeast through homologous recombination, as previously established for fluorescent protein engineering (Knop et al., 2002).

We then evaluated the versatility of both YSD platforms by displaying immunogenic components from three clinically relevant viruses: Severe acute respiratory syndrome coronavirus 2 (SARS-CoV-2), hepatitis C virus (HCV), and hepatitis E virus (HEV). For each viral open reading frame (ORF), regions predicted or reported to be surface-exposed, conformationally stable, and immunogenic were selected for display from literature based on the established domain architecture of the respective proteins (Guu et al., 2009; Lan et al., 2020; Sabo et al., 2011; Wu et al., 2021). Each antigen was fused to either an HA or MYC epitope tag and displayed using the AGA2 platform and/or the GPI platform. To detect surface expression, fluorescent staining of the epitope tag was performed, followed by flow cytometry analysis (Figure 1d). In addition, control strains expressing only the epitope tag (MYC or HA-tag) or lacking any tag (NoTag) were constructed to validate detection specificity.

SARS-CoV-2 antigens displayed robustly on the AGA2 platform, with both HA-tagged and MYC-tagged constructs showing strong surface expression. HEV antigens express very weakly on the AGA2 platform but were successfully presented using the GPI platform, with HA-tagged constructs showing higher expression than the MYC-tagged version, indicating that both the platform and the epitope tag can influence display efficiency. HCV antigens tagged with MYC are also displayed efficiently on the GPI platform (Figure 1e, S1b). These results suggest that antigen display efficiency depends on both the anchoring platform and the epitope tag, and therefore cannot be predicted reliably *a priori*, highlighting the value of a flexible dual-platform YSD strategy.

Another advantage of chromosomally integrated expression constructs is the reduced cell-to-cell variability compared with plasmid-based expression in yeast. We validated the better performance of this system, by using surface display of the SARS-CoV-2 spike receptor-binding domain (S-RBD, amino acids 331 - 531). Comparison of plasmid-based and chromosomally integrated constructs by flow cytometry revealed marked differences in expression homogeneity (Figure 1f). Although both strains expressed the identical HA-tagged AGA2-S-RBD construct, cells carrying the pYD1-derived plasmid exhibited broad heterogeneity in HA fluorescence, whereas the chromosomally integrated strain showed higher and more uniform surface expression. These results demonstrate that genomic integration of the YSD cassette yields more homogeneous protein display than traditional plasmid-based systems, making it well suited for standardized antibody detection applications.

Given the variable display efficiency observed across antigens and platforms, we explored the effect of inserting a robustly folding domain derived from Flo1 (FLO1r; Bester et al., 2006) between the α-factor signal peptide and the linker in the GPI platform. This markedly increased expression of HEV antigens on the GPI platform (Figure 1g). Other optimizations had no obvious effect on display levels (Figure S1c, S1d), whereas alternative signal peptides, such as those from Ost1 and Suc2 (Barrero et al., 2018; Yin et al., 2018), resulted in lower expression on the GPI platform (Figure S1e). We also tested different anchor proteins; however, the stalk-GPI architecture outperformed all alternatives (Figure S1f; Yang et al., 2019). These results identify FLO1r insertion as an effective strategy for enhancing display of challenging antigens and support the α-factor signal peptide and stalk-GPI anchor as optimized components of our platform.

### Quantitative detection of SARS-CoV-2 antibodies using yeast surface display and flow cytometry (YSD-FCM)

We next established a workflow to use YSD strains for quantitative detection of antibodies specific to displayed antigens in human sera (Figure 2a). Since human sera often contain pre-existing antibodies that recognize yeast cell-wall components (Falcon et al., 2025), we introduced a pre-adsorption step to remove yeast-reactive antibodies and minimize nonspecific binding. For this purpose, non-YSD yeast cells lacking both the POI and the epitope tag can be used. For simplicity’s sake, we compared commercially available baker’s yeast with the parental wild-type yeast strain (ESM356-1) used for our YSD system. Commercially available baker’s yeast (fresh baker’s yeast, Wonnemeyer Feinkost) was effective in most cases, but occasionally not sufficient for depletion of all yeast-reactive antibodies (Figure S2a, Supplementary Note 2). This can be explained by the fact that the expressed proteome of individual yeast strains only constitutes a fraction of the entire yeast pan-proteome (Peter et al., 2018), with a substantial fraction of strain-specific proteins at the plasma membrane. Therefore, ESM356-1 cells were used for pre-adsorption in all subsequent experiments unless stated otherwise.

**Figure 2.**
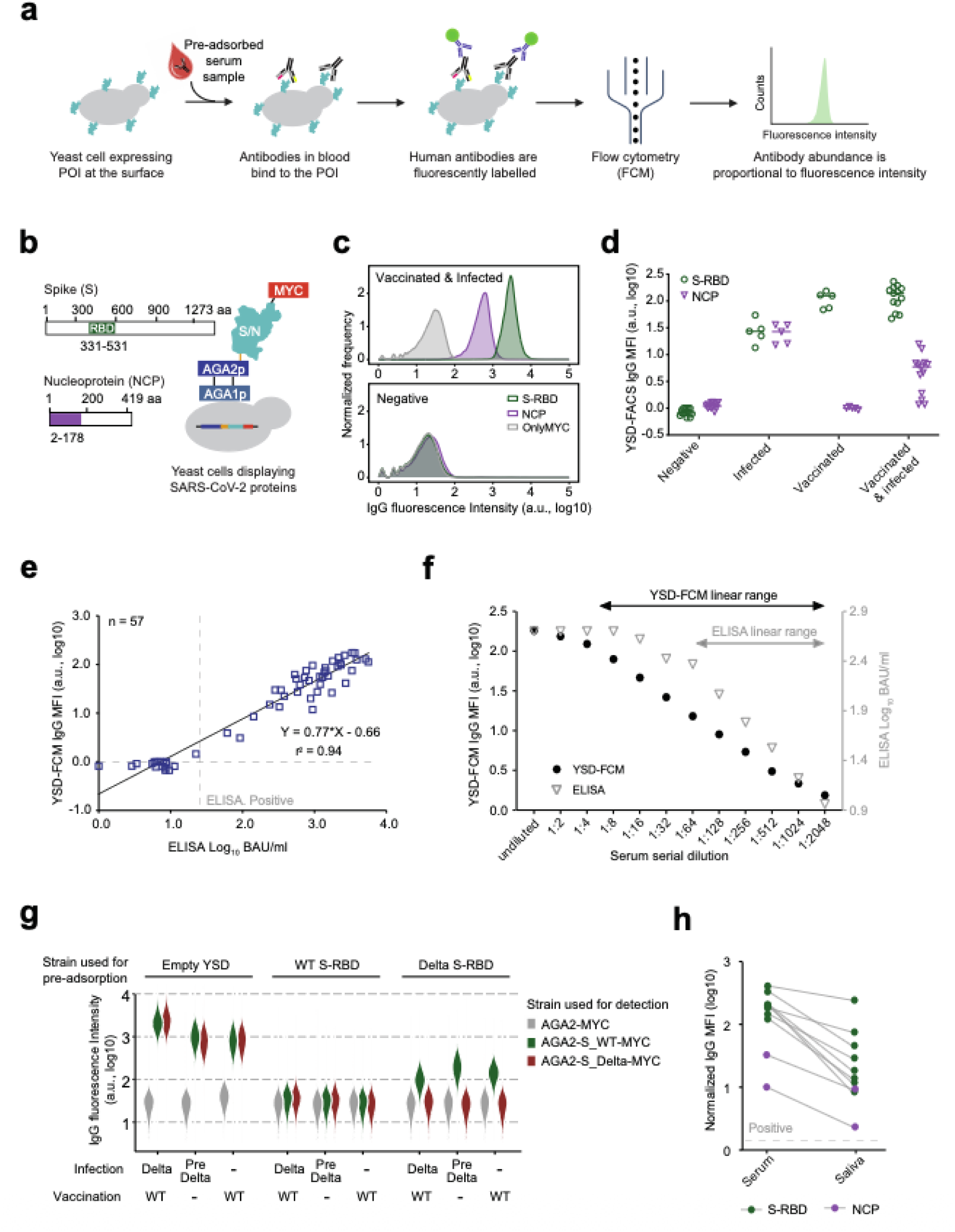
Quantitative detection of SARS-CoV-2 antibodies using yeast surface display and flow cytometry (YSD-FCM). (a) Experimental workflow for quantitative antibody detection in human samples using YSD-FCM. (b) Schematic representation of the SARS-CoV-2 spike (S) and nucleoprotein (NCP), indicating the regions used for yeast surface display. (c) Representative density plots showing antibody detection against S-RBD and NCP in serum from vaccinated-plus-infected donors; negative (pre-COVID-19) serum showed no significant signal. (d) Quantification of anti-S-RBD and anti-NCP antibody levels in sera from vaccinated-only (n=5), infected-only (n=5), vaccinated-plus-infected (n=13), and negative (n=14) individuals. Median fluorescence intensity (MFI; log10-transformed) of human IgG is shown. (e) Correlation of anti-S-RBD antibody titers measured in 57 human serum samples by YSD-FCM and a commercial ELISA. Values on both axes are log-transformed, with YSD-FCM values shown on the y-axis and ELISA titers shown on the x-axis. r2 indicates Pearson’s correlation coefficient. (f) Comparison of the dynamic range of YSD-FCM (black circles, left y-axis) and ELISA (open triangles, right y-axis) using a dilution series of an S-RBD-positive serum sample. (g) Variant-specific antibody depletion using YSD strains displaying wild-type or Delta S-RBD. The violin plot shows human IgG fluorescence intensity measured by flow cytometry, indicating antibody binding to the displayed S-RBD variants after pre-adsorption with different YSD strains. (h) Detection of antibodies against S-RBD or NCP in paired serum and saliva samples using the YSD-FCM assay. Normalized MFI (log10-transformed) of human IgG is shown. The dashed line indicates the positivity threshold for the YSD-FCM signal.

This optimized workflow was then applied to quantify antibodies against SARS-CoV-2 antigens in human sera using the AGA2 platform carrying a C-terminal MYC tag. We first displayed the spike receptor-binding domain (S-RBD), a key immunogenic domain of the spike protein and a major target of antibodies induced by infection and vaccination (Lan et al., 2020; Piccoli et al., 2020; Starr et al., 2020). We also generated a strain displaying a fragment of the nucleoprotein (NCP; amino acids 2–178), which is immunogenic and enables detection of antibodies associated with SARS-CoV-2 infection rather than spike-based vaccination (Figure 2b; i.e. Kadam et al., 2021).

We then analyzed serum from an individual who had previously been infected and vaccinated. This revealed strong fluorescence signals for S-RBD-displaying strains and intermediate signals for NCP-displaying strains, whereas no signal was detected for YSD strains expressing only the AGA2 scaffold protein and carrying the MYC tag. Moreover, pre-COVID-19 sera (negative) showed no significant fluorescence, further supporting assay specificity (Figure 2c). Quantitative analysis of median fluorescence intensity (MFI) allowed clear discrimination of sera obtained from vaccinated-only, infected-only, vaccinated-and-infected, and negative SARS-CoV-2-naïve donors, consistent with their known immunization status (Figure 2d). Fluorescence microscopy further visualized antibody binding on the surface of S-RBD- and NCP-displaying yeast cells, supporting the flow-cytometry results (Figure S2b).

We further investigated parameters influencing the sensitivity of antibody detection. Extending the incubation time of preadsorbed serum with YSD cells to more than 16 hours at 4°C markedly increased fluorescence intensity, particularly for low-titer samples (Figure S2c). The ratio between YSD cell number and serum volume also affected detection sensitivity. Stronger anti-S-RBD antibody signals were observed at lower YSD cell densities, consistent with the idea that anti-S-RBD antibody availability becomes limiting relative to antigen abundance; thus, reducing the number of YSD cells increases the number of antibody molecules bound per cell (Figure S2d). To simplify the workflow and improve reproducibility, we implemented frozen YSD cells for antibody detection, which yielded results highly consistent with those obtained using freshly cultured YSD cells (Figure S2e). Based on these findings, we established a standard operating procedure (SOP; provided upon request) that uses a minimal number of YSD cells per assay, supplemented with wild-type “empty” cells (ESM356-1) as carrier cells to facilitate efficient centrifugation during wash steps. The YSD and carrier cells were premixed and frozen together in large batches, improving reproducibility across experiments.

We then compared the sensitivity and specificity of YSD-FCM with a certified, EU-approved ELISA for SARS-CoV-2 serology, which is based on the S1 domain of the spike protein encompassing the RBD (Beavis et al., 2020). Analysis of 57 human serum samples using both assays demonstrated excellent agreement (*r²* = 0.94) and comparable sensitivity (Figure 2e). Based on the linear regression curve (y = 0.77x - 0.66), YSD-FCM arbitrary units (a.u.) can be correlated to Binding Antibody Units (BAU), which are frequently used in clinical diagnostics. Serial dilution analysis revealed that the YSD-FCM assay maintained a broader linear dynamic range than ELISA, especially at high antibody concentrations (Figure 2f). In addition, replicate measurements of 12 human serum samples for both S-RBD and NCP antibody levels showed high reproducibility, supporting the robustness of the YSD-FCM assay (Figure S2f).

In addition to the MYC-tagged strains, we also generated AGA2-S-RBD and AGA2-NCP strains carrying a C-terminal HA tag to evaluate the impact of the epitope tag on detection performance. The HA-tagged strains showed detection efficiencies comparable to those of the MYC-tagged versions (Figure S2g), indicating that either configuration can support robust surface display and antibody detection. However, in a few serum samples, HA-tagged strains showed elevated background signals due to nonspecific binding (Figure S2h), likely reflecting the presence of antibodies recognizing the HA tag, possibly caused by a prior influenza infection or vaccination.

Beyond quantitative serology, our methodology should also enable epitope-resolved antibody detection, allowing differentiation of immune responses against distinct antigenic variants. To test this, we performed antibody-depletion experiments using YSD strains displaying S-RBD variants from different SARS-CoV-2 strains: Wuhan-Hu-1, referred to here as wild type (WT S-RBD), and Delta virus (B.1.617.2; carrying the Delta-defining RBD substitutions L452R, T478K; referred to as Delta S-RBD). Human sera were obtained from donors who only received the BioNTech mRNA vaccine encoding the Wuhan-Hu-1 spike protein, donors infected with pre-Delta variants, and donors who were vaccinated and subsequently infected with the Delta variant. The sera were preadsorbed with either (i) “empty” YSD yeast cells as a control, (ii) a YSD strain expressing WT S-RBD, or (iii) a YSD strain expressing the Delta S-RBD variant. The preadsorbed sera were then incubated with YSD strains displaying either WT S-RBD or Delta S-RBD.

When “empty” YSD yeast cells were used for pre-adsorption we observed strongest signals for all sera from vaccinated and/or infected individuals when either of the detection strains (WT S-RBD or Delta S-RBD) was used (Figure 2g). Pre-adsorption with WT S-RBD nearly completely eliminated antibody binding in all sera when detected using WT S-RBD. When using the Delta S-RBD YSD strain, we also did not detect antibodies in any of the serum samples from infected and/or vaccinated people, and no signal was detected in the serum derived from an individual that was vaccinated with WT followed by an infection with Delta virus. This could indicate that an infection with the Delta variant (in this WT vaccinated individual) did not promote the formation of detectable amounts of Delta S-RBD specific antibodies. In contrast, pre-adsorption using Delta S-RBD did remove all Delta-specific antibodies, while a significant amount of WT S-RBD specific antibodies were still detected in the sample (Figure 2g, for controls, quantification and microscopic assessment of the data see Figure S3a-c). This indicates that the Delta S-RBD domain lacks binding sites for an important fraction of antibodies generated via immunization or infection with the wild type SARS-CoV-2 strain. Unfortunately, the number of sera with known immunization and infection status was limiting and we did not have access to samples with exclusive Delta virus infection which would be needed to test how many Delta S-RBD specific antibodies are produced in the absence of a vaccination with the WT variant. Therefore these results only provide a preliminary indication that YSD can be used to discriminate against the presence/absence of SARS-CoV-2 lineage specific antibodies.

Finally, we also tested whether the YSD-FCM assay can be used to detect SARS-CoV-2 antibodies in human saliva, a non-invasive and easily obtainable specimen. A total of eight saliva samples, paired with serum samples from the same individuals following SARS-CoV-2 vaccination or infection, were analyzed. S-RBD and NCP antibodies were detected in all positive saliva samples, whereas no signal was observed in negative samples, supporting assay specificity within this small sample set (Figure 2h). Consistent with prior reports (Faustini et al., 2023; Yu et al., 2022), antibody titers in saliva were lower than those measured in serum. Nevertheless, the assay distinguished positive from negative saliva samples in this proof-of-concept analysis, highlighting its potential utility for non-invasive serological surveillance.

### Detection and characterization of sera from convalescent hepatitis E virus patients

To further evaluate the YSD-FCM assay in a distinct clinically relevant viral infection, we applied the workflow to human samples from convalescent patients previously infected with hepatitis E virus (HEV). HEV is a leading cause of acute hepatitis worldwide, with reported seroprevalences of up to 50% in some European cohorts/populations (Lapa et al., 2015). However, currently available commercial HEV serological assays differ substantially in sensitivity, specificity, and estimated seroprevalence, and often show poor concordance between kits, resulting in diagnostic uncertainty (Haller et al., 2025; Zhang et al., 2020). In addition, the growing diversity of *Hepeviridae* members capable of infecting humans - including recently described rat *Rocahepevirus* species (Sridhar et al., 2018) - raises a need for serological tools that can detect divergent HEV-related infections with high specificity (Chen et al., 2025). Together, these challenges highlight the need for high-throughput, adaptable, and specific methods for HEV serology.

The HEV genome encodes three open reading frames: ORF1, which encodes the non-structural replicase machinery, and ORF2 and ORF3, which encodes structural proteins. HEV antigens were expressed using the GPI platform carrying a C-terminal HA tag. First, we generated multiple YSD strains displaying fragments spanning different regions of the ORF2 protein, which together form the capsid protein and represent major targets of the humoral immune response (Figure 3a; Guu et al., 2009). We used the protein sequence of the HEV genotype 3 (HEV-3) strain Kernow-C1, because HEV-3 is the predominant genotype in industrialized countries (Aspinall et al., 2017).

**Figure 3.**
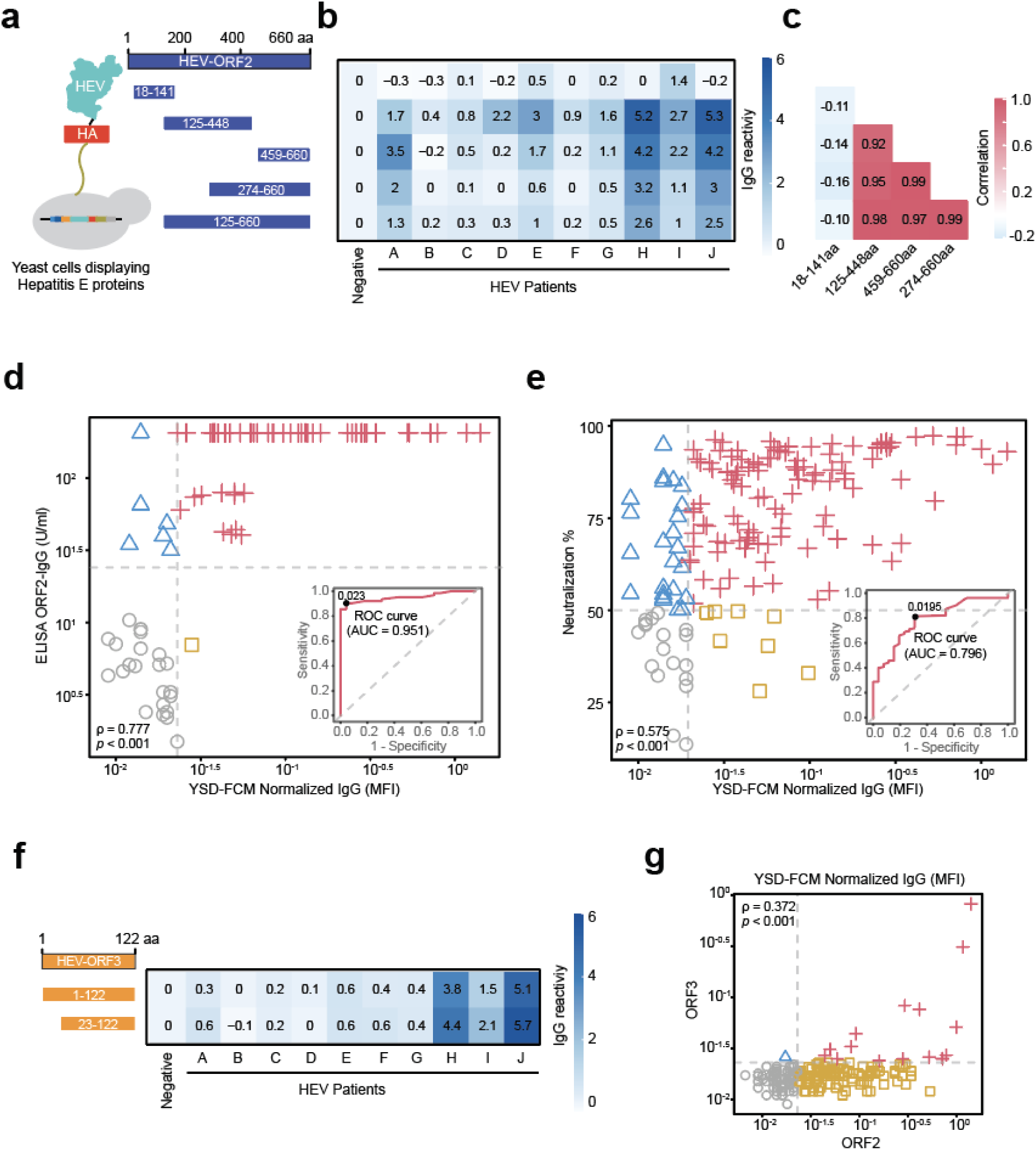
Quantitative detection of hepatitis E virus (HEV) antibodies using yeast surface display and flow cytometry (YSD-FCM). (a) Schematic representation of HEV ORF2 proteins, indicating the regions used for yeast surface display. (b) Heatmap showing the IgG reactivity of ORF2-derived fragments in serum samples from HEV-infected and HEV-negative individuals. IgG reactivity was calculated as the log2 fold change of the IgG/HA ratio relative to the negative serum. (c) Heatmap showing correlation of IgG reactivity among all ORF2-derived fragments. (d) Scatter plot showing correlation between YSD-FCM assay values and ELISA values for 85 serum samples. Spearman’s ρ = 0.777, p < 0.001. Red crosses indicate samples positive by both assays (n = 56); grey circles indicate samples negative by both assays (n = 22); blue triangles indicate samples negative by YSD-FCM assay but positive by ELISA (n = 6); orange squares indicate samples positive by YSD-FCM assay but negative by ELISA (n = 1). The inset (bottom right) shows the Receiver Operating Characteristic (ROC) curve for a binary classifier (red solid line; Area Under the Curve, AUC = 0.951) compared to a random classifier (grey dashed line; AUC = 0.50). The solid black dot on the ROC curve indicates the optimal cutoff threshold (determined by the Youden Index), maximizing sensitivity and specificity. (e) Scatter plot showing correlation between YSD-FCM assay values and neutralization assay values for 161 serum samples. Spearman’s ρ = 0.575, p < 0.001. Red crosses indicate samples positive by both assays (n = 111); grey circles indicate samples negative by both assays (n = 18); blue triangles indicate samples negative by YSD-FCM assay but positive by the neutralization assay (n = 24); orange squares indicate samples positive by YSD-FCM assay but negative by the neutralization assay (n = 8). The inset (bottom right) shows the Receiver Operating Characteristic (ROC) curve for a binary classifier (red solid line; Area Under the Curve, AUC = 0.796) compared to a random classifier (grey dashed line; AUC = 0.50). The solid black dot on the ROC curve indicates the optimal cutoff threshold (determined by the Youden Index), maximizing sensitivity and specificity. (f) Schematic representation of HEV ORF3 proteins, indicating the regions used for yeast surface display. Heatmap showing the IgG reactivity of ORF3-derived fragments in serum samples from HEV-infected and negative individuals. IgG reactivity was calculated as the log2 fold change of the IgG/HA ratio relative to the negative serum. (g) Scatter plot showing ORF2- and ORF3-specific antibody levels in 178 serum samples measured by the YSD-FCM assay. Both axes are shown on a logarithmic scale. Red crosses indicate samples positive for both ORF2-specific and ORF3-specific antibodies (n = 16); grey circles indicate samples negative for both ORF2-specific and ORF3-specific antibodies (n = 64); blue triangles indicate samples positive for ORF2-specific but negative for ORF3-specific antibodies (n = 97); orange squares indicate samples negative for ORF2-specific but positive for ORF3-specific antibodies (n = 1).

Each YSD strain was incubated with serum from 10 convalescent individuals previously infected with HEV or with a negative control serum. Antigen-specific IgG fluorescence was normalized to antigen expression using the HA-tag signal, yielding an IgG/HA ratio. IgG reactivity was then calculated as the log2 fold change in the IgG/HA ratio relative to negative serum. Among the ORF2-derived fragments, all fragments containing sequences that encompass part or the almost full so-called P-domain (“protruding domain”; approx. aa 415 - 710; Guu et al., 2009) showed the strongest IgG reactivity in HEV-positive sera (Figure 3b, S3a). Consistently, this P-domain has been implicated in virus–host interactions and is a major target of neutralizing antibodies (Gu et al., 2015; Ssebyatika et al., 2025). In contrast, amino acids 18-141 showed reactivity close to baseline (Figure 3b, S4a) and correlation analysis showed that the combined IgG responses to fragments 125-448 and 459-660 accounted for most of the ORF2-specific antibody reactivity (Figure 3c). This heterogeneity indicates that the antibody response to ORF2 is not evenly distributed across the protein, but is concentrated on specific regions, likely reflecting differences in epitope availability, accessibility, or antibody affinity. This result demonstrates the use of YSD for mapping immunogenic domains in viral capsid proteins.

We next continued using YSD-FCM for larger-cohort serum screening and comparison with established serological assays. Based on the results in Figure 3c we selected YSD strains displaying fragments aa 125-448 and aa 459-660. The IgG/HA ratios from the two fragments were summed and used as a measure of ORF2-specific antibody levels.

We first evaluated the diagnostic performance of the YSD-FCM assay by comparing it with a widely used ORF2-based ELISA using a set of 85 samples from convalescent HEV patients (Figure S4b; Csernalabics et al., 2024). Spearman’s rank correlation analysis showed a strong positive association between the two assays (ρ = 0.777, *p* < 0.001; Figure 3d). Consistent with this, receiver operating characteristic (ROC) analysis demonstrated strong discriminatory performance, with an area under the curve (AUC) of 0.951 (95% CI: 0.909–0.994). At the selected cutoff, the YSD-FCM assay achieved a sensitivity of 90.3% and a specificity of 95.7% relative to the ELISA test that we used as a reference (Figure 3d). Agreement between the two methods was also high, as indicated by a Cohen’s kappa coefficient of 0.835, consistent with near-perfect agreement. Six samples were classified as negative by YSD-FCM but positive by ELISA, with YSD-FCM values ranging from 0.012 to 0.021 and ELISA values from 1.501 to 2.312. Conversely, one sample was positive by YSD-FCM but negative by ELISA, showing a borderline YSD-FCM signal of 0.028 and a low ELISA value of 0.845 (Figure 3d). These discordant results may reflect differences in antigen format or sequence, assay cutoff placement, or the detection of antibodies against linear versus conformational epitopes. Additionally, genotype variations may contribute to these discrepancies, as the ELISA detects both genotype 1 and genotype 3, whereas this study focuses exclusively on genotype 3.

To determine to what extent the YSD-FCM assay reflects functional neutralizing antibody responses, we compared the YSD results from 161 samples for which neutralizing activity data were available (Csernalabics et al., 2024; Zhang et al., 2022), quantified as percent neutralization at a serum dilution of 1:1,000. Neutralization data were unavailable for the remaining 18 samples (Figure S4b). YSD-FCM signal intensity showed a moderate positive correlation with neutralizing activity (Spearman’s ρ = 0.575, *p* < 0.001; Figure 3e). For ROC analysis, samples with ≥50% neutralization were classified as neutralization-positive, consistent with the common use of 50% inhibitory/neutralization endpoints in virus neutralization assays (Cai et al., 2016; Liu et al., 2019). This analysis showed good discriminatory performance, with an AUC of 0.796 (95% CI: 0.708-0.864). At the selected cutoff, the YSD-FCM assay achieved a sensitivity of 82.2% and a specificity of 69.2% (Figure 3e). Together, these results indicate that although the YSD-FCM assay does not precisely predict the magnitude of neutralization, it can enrich for samples with strong neutralizing antibody activity, supporting its potential use as a rapid screening tool for applications such as vaccine-response assessment, protective immune-status evaluation, and seroprevalence studies.

In addition to ORF2, we designed two ORF3 fragments of different length for use in YSD. ORF3 encodes a short protein that is essential for the release of quasi-enveloped HEV progeny (Yamada et al., 2009). Both fragments showed comparable IgG fluorescence profiles across the tested sera (Figure 3f). Since no commercial reference assay was available for ORF3, direct comparison with an external method was not possible. We therefore screened a total of 178 serum samples using YSD strains displaying the full-length ORF3 fragment (Figure S4b). Compared with ORF2, ORF3-specific antibody responses were generally lower, in line with lower CD4 T-cell responses against ORF3 reported in our previous study (Csernalabics et al., 2024). ORF3-reactive antibodies were detected in 14.2% (16 of 113) of ORF2-positive samples. Although ORF2 and ORF3 antibody levels showed a moderate correlation (Spearman’s ρ = 0.372, *p* < 0.001), their detection was not fully concordant (Figure 3g), suggesting partially overlapping but distinct humoral response patterns. These findings support the inclusion of ORF3 as an auxiliary antigen in future serological panels.

Alongside human HEV, we also generated a small panel of YSD strains expressing the envelope glycoprotein E2 from several hepatitis C virus (HCV) genotypes. E2 was selected because it is a principal target of HCV-specific antibody responses and contains epitopes involved in virus neutralization (Keck et al., 2011; Kong et al., 2015). As a proof of concept, we tested whether these recombinant HCV antigens could be recognized by antibodies present in HCV-infected individuals. A small number of HCV-positive serum samples (n=3) were analyzed against YSD strains displaying E2 fragments derived from genotypes 1a, 1b, and 1c. Robust IgG reactivity was observed for the globally widespread 1a and 1b subtypes when compared with a HCV-negative control serum (Figure S4c). For 1c a weaker response was observed in all 3 samples, which could be explained by cross-reactivity of the sera to this variant that is primarily associated with specific geographic regions (in Africa and Asia) (Messina et al., 2015). These results provide preliminary evidence that the displayed E2 antigens are immunoreactive and suitable for further YSD assay development.

### High-throughput multiplexed antibody profiling using barcoded yeast surface display (SeroSeq)

To scale yeast-surface-display-based serology, we developed SeroSeq, a pooled FACS-sequencing workflow in which each yeast clone displays one antigen and carries a unique DNA barcode (Figure 4a). For each antigen, we generated multiple independently barcoded clones, allowing the same antigen to be included in different pools with distinct barcode identities. Each serum sample was incubated with a separate pool, after which all pools were combined, sorted by FACS according to antibody-binding intensity, and analyzed by barcode sequencing. This design establishes two levels of multiplexing, enabling all POI pools from all serum samples to be analyzed in a single FACS run. (Figure 4a). Thus, SeroSeq converts antibody binding across multiple antigens and serum samples into quantitative barcode-enrichment profiles.

**Figure 4.**
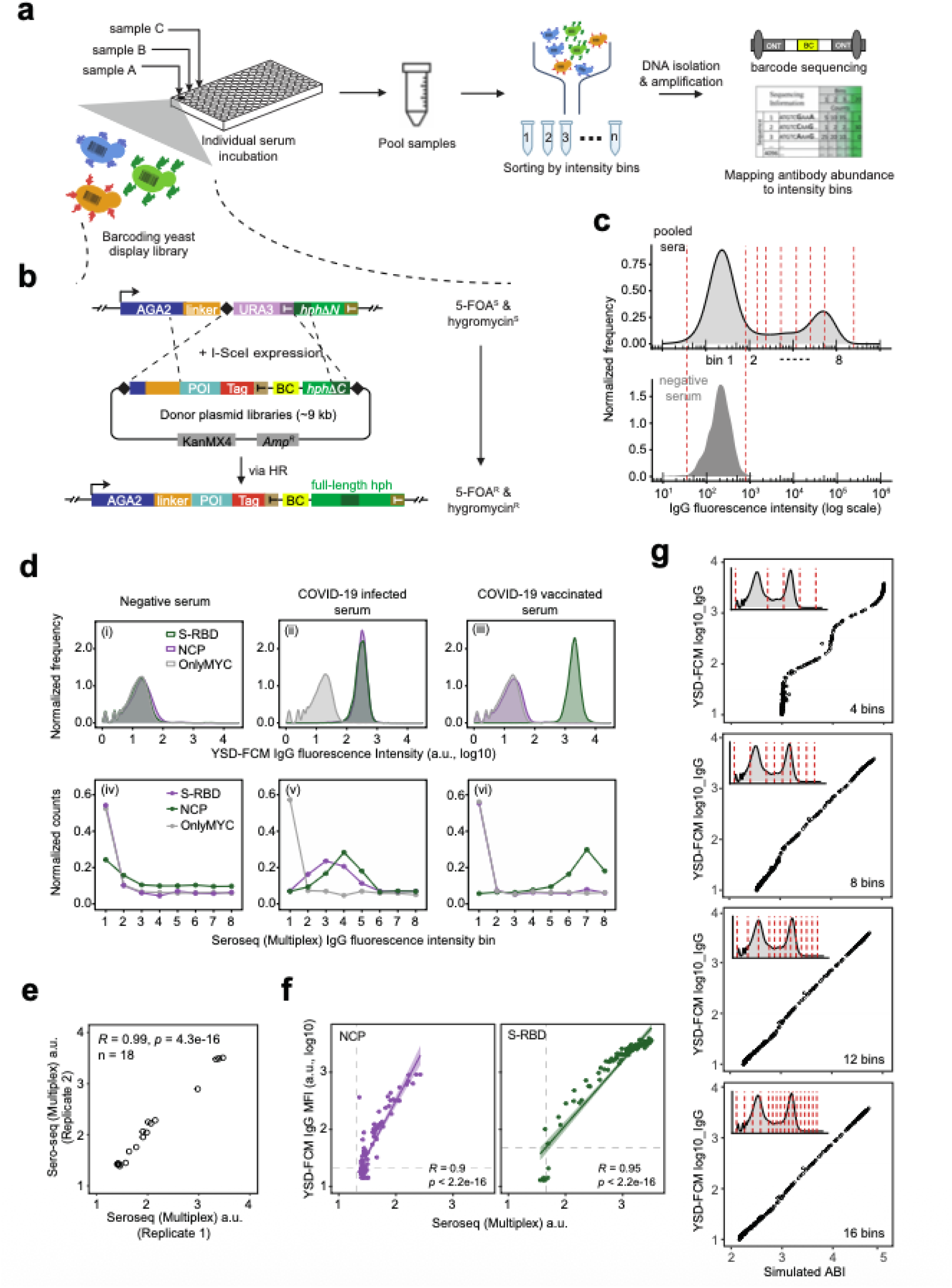
Multiplexed antibody quantification using barcoded YSD libraries and FACS-seq (SeroSeq). (a) Schematic of the SeroSeq workflow. Barcoded YSD strains displaying SARS-CoV-2 S-RBD, NCP, or MYC-tag were mixed in each well and incubated with 96 preadsorbed human serum samples. Cells from all wells were pooled and sorted by FACS, followed by next-generation sequencing. Barcodes recovered from each bin by PCR with bin-specific primers were determined and quantified by high-throughput sequencing. Normalized read distributions across bins were used to infer antibody abundance. (b) Schematic of barcoded YSD library construction strategy using a donor plasmid. Unique barcode oligonucleotides were cloned into donor plasmids encoding the respective POI. After transformation into the AGA2-based acceptor strain, I-SceI induction triggered homologous recombination, integrating a genomically encoded YSD cassette containing the POI and a unique barcode upstream of the reconstituted hphNT1 selection marker. The I-SceI motif is indicated with a black diamond. (c) Binning strategy used for SeroSeq cell sorting. Pooled samples (top panel) were sorted into eight bins according to human IgG fluorescence intensity, with the first bin defined to include the majority of the negative population, using the negative serum (bottom panel) as the reference. Red dashed lines indicate bin boundaries. (d) Representative plots illustrating the intensity profiles of three different serum samples: negative serum, infected serum, and vaccinated serum, as indicated above. The profiles were generated either from single YSD-FCM measurements (panels i–iii) or from SeroSeq followed by analysis of the corresponding sera (panels iv–vi). (e) Duplicate analysis of the SeroSeq assay. Quantification of antibody levels (S-RBD, NCP and MYC) in human serum samples (n=18) using SeroSeq. Identical samples were tested in duplicate using different barcodes. (f) Correlation of antibody levels (S-RBD, NCP, and MYC) determined by single YSD-FCM measurements and SeroSeq (Multiplex) methodology. Of the 96 serum samples initially assayed, 91 passed quality control and were included in the correlation analysis. (g) Evaluation of SeroSeq binning resolution. Single YSD-FCM measurements were computationally partitioned into 4, 8, 12, or 16 bins to simulate SeroSeq sorting strategies. Simulated antibody abundance index (ABI) values calculated from each binning strategy were compared with the corresponding single-sample YSD-FCM measurements.

To validate this strategy, we generated barcoded yeast-display strains presenting SARS-CoV-2 S-RBD, NCP, or a MYC-tag control (Figure 4b, S5a). To this end, we first cloned a pool of unique barcode oligonucleotides into donor plasmids carrying the respective POI. Following transformation of these uniquely barcoded donor plasmids into the AGA2-based acceptor strain, induction of the I-SceI endonuclease triggered in vivo homologous recombination, resulting in genomic integration of a YSD cassette encoding the POI and carrying a unique barcode (BC) upstream of the reconstituted hphNT1 selection marker gene (Figure 4b, S5a). Each clonal strain carried a unique integrated barcode, which was confirmed by sequencing and functionally validated by flow cytometry. All resulting barcoded YSD strains retained MYC expression and IgG fluorescence levels comparable to those of their non-barcoded counterparts, indicating that barcode insertion did not impair surface display or antibody detection (Figure S5b).

We next arrayed the barcoded strains in a 96-well format, with each well containing a defined mixture of one S-RBD strain, one NCP strain, and one MYC-tag strain with known barcodes (Figure 4a). Each preabsorbed human serum sample was incubated with one of these three-in-one YSD mixtures, such that each barcode encoded both the displayed antigen and the corresponding serum sample. The S-RBD and NCP strains were used to detect antigen-specific antibodies, whereas the MYC-tag strain served as a negative control. For antibody staining, all pools were incubated separately with primary and secondary antibodies, followed by washing steps (Figure S6, Supplementary Note 7). We then pooled all cells and sorted them by FACS into eight fluorescence bins, using the negative serum as a reference to define the boundary of the first bin (Figure 4c). Sorted cells were grown to saturation, followed by DNA extraction. Barcodes recovered from each bin by PCR were then identified and quantified by next-generation sequencing. Based on the distribution of normalized read counts across bins, we calculated an antibody abundance index (ABI), which represents the relative antibody-binding signal against each antigen in each sample (Figure 4a).

Barcode read distributions across fluorescence bins revealed distinct serological profiles for negative, infected, and vaccinated sera. In negative serum, S-RBD, NCP, and MYC-tag barcodes were concentrated in the lowest-intensity bin, consistent with background fluorescence. Serum from a SARS-CoV-2-infected individual showed enrichment of both S-RBD and NCP barcodes in intermediate-intensity bins, indicating moderate antibody-binding levels. By contrast, serum from a vaccinated-only individual showed strong enrichment of S-RBD barcodes in high-intensity bins (6–8), whereas NCP barcodes remained largely confined to background levels in bin 1, consistent with spike-specific immunization in the absence of infection (Figure 4d). Duplicate analysis of 18 serum samples showed excellent agreement, supporting the robustness of the SeroSeq workflow (Figure 4e, S7a).

After quality control, 91 of the 96 initially assayed serum samples were retained for quantitative comparison with single-sample YSD-FCM measurements performed on the same serum samples (Figure S7b). ABI values derived from multiplexed SeroSeq showed strong concordance with the corresponding single-sample YSD-FCM measurements, demonstrating the quantitative performance of SeroSeq (Figure 4f, S7c). Concordance was reduced for negative and weakly positive samples, likely because the broad first bin limited resolution in the low-fluorescence range (see Discussion).

We realized that the binning strategy used in this experiment was not optimal, particularly for negative and weakly positive samples. To optimize the quantitative resolution of SeroSeq, we computationally evaluated alternative binning strategies using single-cell measurements obtained from individual YSD-FCM assays of 96 serum samples against SARS-CoV-2 S-RBD, NCP, or MYC-tag strains (Figure S7d). The combined single-cell dataset was divided into 4, 8, 12, or 16 fluorescence-intensity bins to simulate SeroSeq sorting, and the resulting simulated ABI values were compared with the corresponding single-sample YSD-FCM measurements (Figure 4g). Using only four bins resulted in a compressed, step-like distribution with poor resolution, particularly among negative and weakly positive samples. Increasing the number of bins to eight, by subdividing each bin from the four-bin strategy into two bins, improved overall agreement but retained some discretization. In contrast, the 12- and 16-bin strategies produced near-linear concordance across the full dynamic range, including the low-intensity region (Figure 4g). These results indicate that increasing the number of sorting bins improves the quantitative resolution of SeroSeq measurements. Although 12 or 16 bins provided the highest resolution, the eight-bin strategy represented a practical balance among quantitative performance, sorting time, and experimental complexity. Given the clear outcome of the computational analysis and the costs associated with sequencing, we did not perform an additional experimental validation.

Together, these results establish SeroSeq as a robust and quantitative high-throughput serological profiling platform, while identifying bin number as a key parameter governing sensitivity in the low-titer range.

## Discussion

Accurate quantification of antigen-specific antibodies is essential for infectious disease diagnostics, vaccine evaluation, seroepidemiology, and immunomonitoring (Crescioli et al., 2026; Long et al., 2020; Lu et al., 2018; Paul et al., 2024; Schroeder & Cavacini, 2010). However, many serological platforms remain limited in their ability to combine quantitative performance, antigen flexibility, scalability, and straightforward assay standardization. Here, we established a chromosomally integrated yeast surface display platform that combines rapid strain construction, stable antigen presentation, flow-cytometric antibody quantification, and barcode-enabled multiplexing. Across viral antigens from SARS-CoV-2, HEV, and HCV, the platform enabled robust and reproducible detection of antigen-specific antibodies in human samples, supporting its utility as a flexible framework for serological assay development and immune profiling. In addition, yeast is a renewable and readily scalable resource that can be expanded rapidly and cost-effectively when demand increases, for example during a pandemic.

A central feature of the system is its modular display architecture. By implementing both Aga2- and GPI-anchor-based formats, we accommodated antigens with different structural and biochemical requirements. Display efficiency varied substantially between antigens, anchoring strategies, and epitope tags, as illustrated by HEV antigens that displayed poorly in the Aga2 platform but robustly in the GPI platform. These findings highlight that optimal antigen presentation cannot be reliably predicted a priori and support the parallel testing of display configurations when establishing new serological targets. Chromosomal integration further improved the system by reducing expression heterogeneity compared with plasmid-based YSD, thereby providing more uniform antigen presentation and improving the basis for quantitative measurements. Together with epitope-tag-based normalization and defined negative controls, this design supports robust and reproducible antibody detection.

YSD-FCM showed strong analytical performance in comparison with established serological assays. For SARS-CoV-2, the assay showed excellent agreement with a commercial ELISA and provided a broader linear dynamic range, particularly at high antibody concentrations. The workflow was further facilitated by effective removal of nonspecific yeast-reactive antibodies through pre-adsorption, and by the use of frozen YSD cell preparations, which simplified assay handling and standardization. Consistent results across replicate measurements further confirmed the reproducibility and robustness of the assay. For HEV, YSD-FCM strongly correlated with an ORF2-based ELISA and achieved high diagnostic sensitivity and specificity. In addition, analysis of multiple ORF2 fragments revealed that antibody reactivity is concentrated in discrete immunodominant regions, demonstrating the value of YSD for mapping antigen-specific humoral responses within a common experimental framework.

The comparison with HEV neutralization data indicates that YSD-FCM captures antibodies directed against functionally relevant epitopes, although it does not replace direct neutralization assays. The moderate correlation between binding signal and neutralizing activity is expected, because total binding antibodies and neutralizing antibodies only partially overlap. Nevertheless, the assay identified samples with strong neutralizing activity, suggesting potential utility as a screening tool when full neutralization testing is impractical. Beyond total antibody quantification, variant-specific depletion experiments with SARS-CoV-2 S-RBD further show that YSD-FCM can resolve antigenic specificity at the level of related antigen variants. This capability may be useful for dissecting antibody responses to viral evolution, vaccination, breakthrough infection, or other settings in which epitope selectivity is biologically or clinically informative.

To extend the platform toward higher-throughput profiling, we developed SeroSeq, a pooled FACS-sequencing workflow based on barcoded YSD strains. In this format, DNA barcodes encode antigen and sample identity, enabling multiplexed analysis of multiple sera and antigens in a single FACS experiment. In a proof-of-concept SARS-CoV-2 application, SeroSeq-derived antibody abundance indices showed strong concordance with matched single-sample YSD-FCM measurements, demonstrating that barcode-based pooling can preserve quantitative performance while increasing throughput. Computational analysis further identified bin number as an important determinant of quantitative resolution, particularly for negative and weakly positive samples. Increasing the number of fluorescence bins improved agreement across the dynamic range, whereas an eight-bin strategy provided a practical compromise between resolution, sorting time, sequencing cost, and experimental complexity.

Several limitations remain. First, although the platform was validated using antigens from three viruses and applied to several human sample cohorts, broader validation across additional pathogens, autoimmune antigens, and independent clinical cohorts will be required to define its diagnostic scope. Second, antigen format remains an important variable, and differences between YSD-FCM, ELISA, and neutralization assays may reflect distinct presentation of conformational, linear, denatured, or strain-specific epitopes. Third, the current SeroSeq implementation has reduced resolution in very low-titer samples, which could be improved through optimized binning, alternative scoring strategies, or increased sequencing depth. Finally, some biologically informative sample groups, such as sera from individuals infected only with specific SARS-CoV-2 variants, were not available and will be required for more complete evaluation of variant-specific antibody profiling.

Several plasmid-based yeast surface display approaches have previously been used for antibody discovery, epitope mapping, and serum antibody profiling. In these studies, YSD has enabled the selection and engineering of antibody fragments, the discovery of conformationally selective nanobodies, the mapping of monoclonal and polyclonal antibody escape across SARS-CoV-2 RBD variant libraries, and the identification of autoantibody targets using barcoded exoproteome libraries. These applications demonstrate the power of YSD for interrogating antibody–antigen interactions at high throughput. However, their primary focus has generally been antibody discovery, antigen discovery, or mechanistic mapping of immune escape rather than standardized quantitative serology.

Our platform extends this concept toward diagnostic and immune-monitoring applications. By using chromosomally integrated YSD constructs, we reduce variability associated with plasmid copy number and improve the uniformity of antigen display, providing a more robust basis for quantitative antibody measurement. In addition, SeroSeq combines stable antigen display with barcode-based sample and antigen encoding, FACS-based separation by antibody-binding intensity, and sequencing-based quantification. This enables multiple antigens and serum samples to be analyzed in parallel while preserving a quantitative fluorescence-linked readout. Thus, in contrast to previous plasmid-based YSD approaches that primarily identify antibody targets or escape variants, SeroSeq is designed to provide scalable, quantitative serological profiling suitable for diagnostic assay development, serosurveillance, and immune monitoring.

Several previous studies have explored yeast surface display as a format for antibody detection and serological profiling. Early work established yeast-displayed antigens as whole-cell immunoassay reagents, including cell-counting and microfluidic formats for multiplex antibody detection (Guo et al., 2010; Wang et al., 2012). More recently, YSD-based serological assays have been used to profile human IgG binding to SARS-CoV-2 RBD variants (Lopez-Morales et al., 2023). In parallel, barcoded yeast-displayed antigen libraries have been used for high-throughput immune profiling, including mapping of polyclonal plasma antibody escape from SARS-CoV-2 RBD variants and discovery of autoantibodies against the human exoproteome (Greaney et al., 2021a, 2021b; Wang et al., 2022). These studies demonstrate the value of YSD for presenting folded antigens and interrogating antibody specificity at high throughput. However, most of these approaches were designed primarily for immune-escape mapping, antigen discovery, or target identification rather than standardized quantitative serology.

In summary, we present a chromosomally integrated YSD platform for flexible, quantitative, and scalable antibody detection in human samples suitable for diagnostics. The system combines stable antigen display, single-cell normalization, adaptable antigen presentation, and compatibility with pooled barcode-based readout. Its performance relative to established serological assays, broad dynamic range, and ability to interrogate antigen-specific, variant-specific, and multiplexed antibody responses support its use for serological assay development, immune monitoring, and high-throughput antibody profiling.

## Methods

### Yeast methods

All strains were constructed using recombinant DNA (plasmids or linear DNA fragments) using frozen competent cells and LiAcetat yeast transformation, as described previously (Knop et al., 1999), using principles of good microbial practice, as described in Guthrie and Fink, (1991).

### Yeast surface display strain construction

All strains used in this study are listed in Table S1. Most strains were derived from *Saccharomyces cerevisiae* ESM356-1 or ESM357-9 (Knop & Schiebel, 1998), both spores from strain FY1679 (Winston et al., 1995).

### Acceptor strains for AGA2-based chromosomally integrated yeast surface display

For AGA2-based yeast surface display (YSD), four acceptor strains were used: HHY8699, HHY8631, HHY8397, and HHY8465. Each strain contained a distinct version of the acceptor module. Over the course of the project, the acceptor module underwent several rounds of modification, testing, and refinement to address practical considerations related to cloning and strain construction. This iterative process generated multiple versions of the AGA2-based acceptor strain. Of these, HHY8699 represents the final optimized version and is recommended for future applications.

**HHY8699** was generated from plasmid pHHB1468 using primers HHP4744 and HHP4874. Its acceptor module contains pGPD-AGA2, the *S. paradoxus CYC1* terminator, I-SceI followed by URA3, and an N-terminally truncated hphΔN fragment corresponding to amino acids 146–342, followed by the *S. paradoxus ALG9* terminator (Fig. 1a(iii), Fig. S1a, Table S1).

**HHY8631** was generated from plasmid pHHB1459 using primers HHP4744 and HHP4745. Its acceptor module contains pGPD-AGA2, I-SceI followed by URA3, and an N-terminally truncated hphΔN fragment corresponding to amino acids 146–342, followed by the *S. paradoxus ALG9* terminator. Unlike HHY8699, this module lacks the *S. paradoxus CYC1*terminator (Fig. S1a, Table S1).

**HHY8397** was generated from plasmid pHHB1386 using primers HHP4790 and HHP4745. Its acceptor module contains a C-terminally truncated hphΔC fragment corresponding to amino acids 1–192, pGPD-AGA2, URA3 followed by TrxA and I-SceI (Fig. S1a, Table S1).

**HHY8465** was generated from plasmid pHHB1393 using primers HHP4790 and HHP4745. Its acceptor module contains a C-terminally truncated hphΔC fragment corresponding to amino acids 1–192, pGAL1-AGA2, URA3 followed by TrxA and I-SceI (Fig. S1a, Table S1).

The amplified acceptor module was then integrated at the *HOM3* locus by homologous recombination (selection on hygromycin plates). Correct genomic integration was verified by diagnostic PCR and Sanger sequencing.

### Acceptor strains for GPI-based chromosomally integrated yeast surface display

The acceptor strains used to generate chromosomally integrated, GPI-anchored YSD strains were HHY8851, HHY8360, and HHY8240. These strains differed in the configuration of the truncated hph cassette (*hphΔC* versus *hphΔN*), the epitope tag (HA versus MYC), and the presence or absence of the (GGGGS)₃ linker. HHY8851 represents the final version of the GPI-based acceptor strain and is recommended for future applications of this method.

**HHY8851** was generated from plasmid pHHB1501 using primers HHP4952 and HHP4745. Its acceptor module contains an N-terminally truncated *hphΔN* fragment corresponding to amino acids 146–342, followed by the *S. paradoxus ALG9*terminator, *pGPD*-*MFα*, a (GGGGS)₃ linker, *URA3* with its endogenous promoter and terminator, an I-SceI endonuclease recognition site, a MYC tag, a 649-amino-acid stalk sequence, and a C-terminal glycosylphosphatidylinositol (GPI) anchor sequence (Fig. 1b(iv), Fig. S1a, Table S1).

**HHY8360** was generated from plasmid pHHB1375 using primers HHP4790 and HHP4745. Its acceptor module contains a C-terminally truncated *hphΔC* fragment corresponding to amino acids 1–192, *pGPD*-*MFα*, a (GGGGS)₃ linker, *URA3* with its endogenous promoter and terminator, an I-SceI endonuclease recognition site, an HA tag, a 649-amino-acid stalk sequence, and a C-terminal GPI anchor sequence (Fig. 1b(iv), Fig. S1a, Table S1).

**HHY8240** was generated from plasmid pHHB1348 using primers HHP4744 and HHP4745. Its acceptor module contains *pGPD*-*MFα*, *URA3* with its endogenous promoter and terminator, an I-SceI endonuclease recognition site, an HA tag, a 649-amino-acid stalk sequence, and a C-terminal GPI anchor sequence. Unlike HHY8851 and HHY8360, this module lacks the (GGGGS)₃ linker and truncated hph cassette (Fig. 1b(iv), Fig. S1a, Table S1).

The amplified acceptor modules were integrated at the *HOM3* locus by homologous recombination. Correct genomic integration was verified by diagnostic PCR and Sanger sequencing.

### Design and generation of antigen-encoding DNA fragments

The protein-of-interest (POI) sequences displayed on the yeast surface were designed based on published information and/or predicted structural features. Selected regions were chosen (more or less empirically) to correspond to putative stable folding units, thereby increasing the likelihood of proper folding, escape from cellular quality-control pathways, and efficient surface exposure. DNA fragments encoding the selected regions were obtained either by PCR amplification from existing template plasmids or by de novo synthesis as double-stranded DNA gBlocks (Integrated DNA Technologies or equivalent). Unless otherwise specified, all S-RBD constructs used in this study correspond to the wild-type Wuhan-Hu-1 sequence. The plasmids and gBlocks used in this study are listed in Table S2.

### Donor plasmids for construction of chromosomally integrated yeast surface display acceptor strains

Plasmids used for integration of the acceptor modules are listed in Table S2. For cloning, standard cloning procedures and synthetic DNA were used. Annotated plasmid files are available upon request.

### Integration of POIs in YSD acceptor strains

To generate chromosomally integrated YSD strains using the two-fragment co-transformation approach, I-SceI expression was induced with galactose for 6-24 hours prior to harvesting the cell for yeast transformation. Two DNA fragments carrying the POI and repair elements, respectively, were transformed into the acceptor strains (Fig. 1a). Transformants were selected on Synthetic complete plates containing monosodium glutamate as N-source, (SC-MSG), supplemented with 5-fluoroorotic acid (5-FOA; 1 g/L) and hygromycin B Gold (200 mg/L; InvivoGen).

For POI integration using the plasmid-based donor approach, the donor plasmids carrying the POI were transformed into the acceptor strains (Fig. 1a), and transformants were selected on YPD medium supplemented with G418 (200 mg/L). Single colonies were inoculated into SC-Gal/Raff medium, consisting of synthetic complete medium containing 2% (w/v) raffinose and 2% (w/v) galactose, to induce I-SceI cleavage. Cells were subsequently plated on SC-MSG medium supplemented with 5’-FOA and hygromycin for selection of chromosomal integrants. Correct genomic integration was verified by diagnostic PCR and Sanger sequencing.

### Generation of plasmid-based YSD strains

The plasmid-based YSD strain used in Figure 1f was constructed by transforming plasmid pHHB1380 into strain HHY8151, which overexpresses the *AGA1* gene under the control of the inducible *GAL1* promoter. Transformants were selected on SC-Trp plates. Plasmid-based YSD strains with GPI platform were used to investigate the factors affecting surface display efficiency (Figure 1g, S1d-S1f). These strains were generated by transforming plasmids carrying the corresponding POI fragments into the wild-type strain ESM356-1, followed by selection on SC-Trp plates.

The used plasmids are listed in Table S2.

### Generation of barcoded donor plasmid libraries and yeast libraries

A 69-bp oligo library containing a central 30-nt random barcode was amplified with primers HHP4869/HHP4870 using ten parallel PCR reactions to reduce amplification bias. The pooler PCR product was purified, digested with EcoRI/XmaI restriction enzymes, and cloned into donor plasmids pHHB1485, pHHB1489, and pHHB1525. Each plasmid library yielded at least 1.8 × 10⁴ transformants, which were pooled for plasmid isolation and transformed into yeast acceptor strain HHY8699. Following G418 selection, at least 120 transformants per library were induced for I-SceI-mediated recombination and selected on 5-FOA/hygromycin. Barcode regions were PCR-amplified, verified by Sanger sequencing, and 96 validated strains were arrayed in 96-well format for storage at −80°C. Plasmids and oligos are listed in Tables S2 and S3.

Validated MYC-tag, S-RBD, and NCP YSD strains were expanded from 96-well stocks, normalized by cell density, and mixed with the empty carrier strain ESM356-1. For each well, 3.15 × 10⁶ ESM356-1 cells were mixed with 4.5 × 105 cells of each barcoded YSD strain, resulting in a final mixture of 70% carrier cells and 10% each YSD strain. Plates were centrifuged, supernatants removed, and stored at −80°C until antibody quantification.

### Quantification of YSD surface expression and human antibodies by flow cytometry (YSD-FCM)

YSD cells were grown to logarithmic phase in SC medium (or SC-Trp medium for plasmid-based YSD cells), harvested and subjected to immunostaining depending on the epitope tag (MYC or HA).

For surface expression analysis, cells were incubated with rat anti-HA (Merck, 11867423001) or mouse anti-MYC antibody (Merck, 05-724MG) diluted 1:200 for 30 min at room temperature, washed twice, and stained with Alexa 594-conjugated anti-rat (Invitrogen, A21471) or anti-mouse (Invitrogen, A11032) secondary antibody diluted 1:100 for 30 min at room temperature. After two final washes, cells were resuspended in 200 µl 1x TBS and analyzed by flow cytometry.

For human antibody quantification, human samples (blood, serum, plasma, or saliva samples) were preadsorbed with non-YSD yeast cells, including commercial baking yeast, ESM356-1, or scaffold-only AGA2/GPI strains lacking the tag and POI. In brief, 2.5 µl sample was incubated with 5×10^8^ non-YSD yeast cells in 250 µl 1x TBS containing anti-MYC or anti-HA antibody for 2 hours at 4°C. After centrifugation, 50 µl of the preadsorbed supernatant was transferred to YSD strains and incubated overnight at 4°C with rotation. Cells were washed and stained for 30 min at 4°C with anti-human IgG-DyLight488 (Invitrogen, SA5-10126) and anti-mouse H+L-Alexa 594 (Invitrogen, A11032) (each diluted 1:100), washed again, and resuspended in 150 µl TBS for analysis.

Flow cytometry was performed on a BD FACS Canto^TM^ cytometer equipped for detection of both green and red fluorescence channels. Rainbow Calibration particles (Sphero^TM^, 556286, BD Biosciences) were included in every experiment for calibration/normalization purposes.

### Fluorescence microscopy

Cells were grown in 5 ml SC medium overnight. Next morning cells were diluted to OD_600_ = 0.2 in 20 ml SC medium and grown for 5 hours. Yeast cells were incubated for 30 min at room temperature with preadsorbed seras (and anti-MYC antibody) according to protocol described above. Next, cells were washed incubated for 30 min at 4°C with anti-human IgG-DyLight 488 (Invitrogen, SA5-10126) and anti-mouse H+L-Alexa 594 (Invitrogen, A11032) secondary antibodies, both used at a 1:100 dilution in 1x TBS. Cells were washed 3 times with 1x TBS, and then attached to glass-bottom 96-well microscopy plates (MGB096-1-2-LG-L, MatriPlate) using Bioconext/Concanavalin A coating, as described previously (Khmelinskii & Knop, 2014). Images were taken with a Nikon Ti-E epifluorescence microscope equipped with a 60x ApoTIRF oil-immersed objective (1.49 NA, Nikon), a 2048 x 2048 pixel (6.5 μm), an sCMOS camera (Flash4, Hamamatsu), and an autofocus system (Perfect Focus System, Nikon) with bright field, 469/35 excitation and 525/50 emission filters, or 542/27 excitation and 600/52 emission filters (all from Semrock, with exception of 525/50 which is from Chroma).

### Enzyme-linked immunosorbent assays (ELISA)

SARS-CoV-2 S-RBD-specific IgG antibodies were quantified using the QuantiVac ELISA kit (Euroimmun, EI 2606-9601-10). SARS-CoV-2 NCP-specific IgG antibodies were semi-quantitatively measured using the NCP ELISA kit (Euroimmun, EI 2606-9601-2). IgG antibodies against Hepatitis E Virus (HEV) were quantified using the recomWell HEV IgG ELISA kit (Mikrogen 5004). Human serum samples, calibrators, and controls were processed according to the manufacturers’ instructions. Absorbance was measured at 450nm using a plate reader (Tecan Spark Cyto). ELISA results were interpreted according to the respective manufacturer’s cutoff.

### Multiplexed quantification of antibodies using fluorescence-activated cell sorting and sequencing analysis (SeroSeq)

The multiplexed quantification of antibodies was performed as described previously for YSD-FCM-quantification with the following modifications. After the third wash step of the secondary antibodies, all the samples were pooled and washed one more time with 14.4 ml 1x TBS. Cells were centrifuged and the yeast pellet was resuspended in 4 ml TBS for further fluorescence - activated cell sorting (FACS) using a BD FACS Aria^TM^ III Cell Sorter equipped for the detection of green fluorescent proteins (excitation: 488nm, long pass 502, bandpass: 530/30) and red-fluorescent proteins (excitation: 561nm, long pass 600, bandpass: 610/20).

A total of 2×10^6^ MYC+ (Alexa594) cells were sorted in 8 fractions (bins) according to the log_10_-transformed fluorescence emission intensity on the DyLight488 channel (anti-human IgG signal). The number of sorted cells per bin was adjusted according to the percentage of cells found in each bin. Sorted cells were grown overnight in YPD medium and then harvested for genomic DNA isolation.

### Genomic DNA isolation

Yeast cells were resuspended in 500 µl S-buffer (10 mM K_2_HPO_4_ pH 7.2, 10 mM EDTA, 50 mM β-mercaptoethanol) and incubated with zymolyase 100T at 37 °C for 30 min. Then, 100 µl lysis buffers (25 mM Tris-HCl pH 7.5, 25 mM EDTA, 2.5 % (w/v) SDS) were added, vortexed and incubated at 65°C for 30 min. Proteins were precipitated by addition of 166 µl 3 M potassium acetate, followed by 10 min incubation on ice and centrifugation for 10 min at 4 °C 14,000 rpm. DNA was precipitated from the supernatant by addition of 800 µl cold 100% ethanol and centrifugation. The DNA pellet was washed with 500 µl 70 % ethanol (v/v) and resuspended in 40 µl dH_2_O. Next, the DNA was treated with 1 µl RNase A 10 mg/ml at 37°C for 30 min, followed by addition of 800 µl 70 % EtOH and centrifugation. The DNA pellet was resuspended in 25-50 µl dH_2_O.

### Amplification of barcodes and Oxford Nanopore sequencing

The DNA obtained from each bin (pool of cells) was used as a template for PCR amplification. A 450 bp genomic region containing a 30-mer barcode (unique for each antigen and position per well) was amplified using NEB LongAmp Hot Start 2x master mix (NEB M0533S) and primers containing additional barcodes used to distinguish bins 1-8. A list containing primers used for the barcoding (in addition to the ones in Nanopore PCR kit Oxford Nanopore Technologies, SQK-PSK004) are indicated in Table S3. PCR reactions were conducted using 100 ng genomic DNA, and the following PCR settings: 94 °C 1 min denaturation, followed by 25 cycles: 94°C 30 sec denaturation, 55 °C 30 sec annealing, 65 °C 30 sec extension, followed by 65 °C 5 min. An equivalent amount of PCR products for bins 1-8 were pooled, and 50 µl of this pooled PCR was purified using AMPure XP magnetic beads.

Purified DNA was quantified with Qubit fluorometer and 30 ng were used for a second PCR reaction using Nanopore PCR kit (SQK-PSK004). The following PCR program was used: 94 °C 1 min denaturation, followed by 30 cycles: 94 °C 30 sec denaturation, 62 °C 30 sec annealing, 65 °C 30 sec extension, followed by 65 °C 5 min. The PCR was purified with AMPure XP magnetic beads and 100 fmol of the PCR product were used for sequencing library preparation according to manufacturer’s protocol. The DNA library was loaded on a MinION Mk1b device with a SpotON R9.4.1 Flow Cell.

### Analysis of Nanopore sequencing data and read counting

Raw signal data were generated in FAST5 formats and basecalled and demultiplexed using Guppy v6.2.1 (ONT). Each DNA fragment analyzed by Nanopore sequencing contained three unique barcodes: one internal barcode (30N) identifying the antigen and the position in the 96 - well plate, and two flanking barcodes (24N) that defined the fluorescence-intensity bin from the FACS sorting. Sequencing reads were processed using custom R scripts to extract informative regions by identifying the internal barcode and at least one of the distal barcodes, thereby assigning each read to its corresponding antigen and fluorescence-intensity bin. Read counts were normalized to the total number of reads in each bin, and an antibody abundance index (ABI) was calculated to quantify overall antibody abundance. The ABI was defined as:

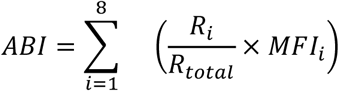

where *R_i_* is the read count for bin *i*, *R_total_* is the total number of reads across all bins, and *MFI_i_* represents the mean fluorescence intensity of bin *i* obtained from FACS data.

### Statistical analysis

For the detection of SARS-CoV-2 antibodies, correlation between the YSD-FCM assay and ELISA was assessed using Pearson’s linear correlation, performed by Prism (version 10.2.3). For the detection of HEV antibodies, correlation between the YSD-FCM assay and ELISA, Neutralization assay, and the correlation between HEV-ORF2 and HEV-ORF3 within YSD-FCM assay was assessed using Spearman’s rank correlation coefficient due to non-normal distribution of values (Shapiro-Wilk test). ROC curve was generated to evaluate the diagnostic performance of the YSD-FCM assay relative to ELISA and neutralization assay. ELISA IgG values or neutralization activity values were used as the reference standards, and the normalized IgG fluorescence signals from the YSD-FCM assay were used as the predictor variable. The area under the ROC curve (AUC) was calculated with 95% confidence intervals (CI). Receiver operating characteristic (ROC) analysis was performed using R (version 4.5.1) with the *pROC* package, and balanced thresholds (optimal cutoffs) were determined using Youden’s J statistic (maximizes sensitivity + specificity - 1) for general screening (Robin et al., 2011). Categorical agreement was evaluated using Cohen’s kappa coefficient (κ), which quantifies concordance between categorical outcomes while correcting for agreement expected by chance (Landis and Koch, 1977). All statistical analyses were performed using either R or Graphpad Prism.

## Ethics approval

The use of samples was covered by the respective ethics approvals and consent procedures of the original studies and sample collections. The studies involving samples approved in Heidelberg were reviewed and approved by the Ethical Review Board of the Medical Faculty of Heidelberg University Hospital, Heidelberg University, Germany (approval number S-200/2021 and S-148/2020). One additional study was approved by the Ethics Committee of the Albert-Ludwigs-University Freiburg, Germany (approval numbers 474/14, 201/17, and 486/19). Written informed consent was obtained from all blood donors prior to enrollment in the respective studies.

Existing samples were re-analyzed by applying our yeast surface display-based serological method to repeat, extend and validate previously performed ELISA-based serological measurements. The relevant ethics approvals and informed consent documents included permission for additional serological analyses of the collected samples. All procedures were conducted in accordance with the applicable ethical standards and relevant guidelines.

The study was conducted in accordance with the Declaration of Helsinki ethical standards. Participants provided consent before sample and data collection.

## Author contributions

DL, HH, MK and VLDT designed research; DL, HH, AD, LM, KH, HYL, MM, DK, IM, and EG designed and performed the experiments; DL, HH, FH, LM, KH, VLDT, and MK analyzed and interpreted data; DL, HH, and MK wrote the manuscript with input from all authors; MK and EG secured funding; IM, PS, UM, CD, TB and VLDT provided technical or material support; MK supervised the research. All authors reviewed and approved the manuscript.

## Data availability

Data are available upon request to the corresponding author.

## Acknowledgements

We would like to thank current and former members of the lab, members of the Heidelberg “Coronatest team” as well as colleagues and friends of our labs for supporting this study. In particular, we thank Robin Burk, Krisztina Gubicza, Christian Reinbold, Gautier Debaeker, Elena Chassis du Guerny, Heike Alter, Andreas Deckert, Michael Persike, Maria Gomez-Fabra Gala, Nikolai Gunkel, and Ioana Huber. We are equally grateful to the many additional colleagues and friends who supported this study in various ways and who preferred not to be acknowledged by name. We thank Monika Langlotz and the ZMBH FACS facility for technical assistance and support. Parts of the test development and validation were supported by the Baden-Wuerttemberg Ministry of Science, Research and Art, as well as by funding received within the framework of the Excellence Strategy of the Federal and State Governments of Germany embedded in the nationwide research network “Applied Surveillance and Testing” (Bundesweites Forschungsnetz “Angewandte Surveillance und Testing”, B-FAST) and the University Medicine’s Network (Netzwerk Universitätsmedizin, NUM) for COVID-19 research. This study was further supported by CRC/TRR179 project 04 (to T.B.), project 22 (to V.L.D.T) of the German Research Foundation (DFG; no. 272983813) and the Carl-Zeiss-Stiftung (to K.H. as part of Center SynGen). Claudia Denkinger is supported through the Heisenberg Program of the German Research Foundation (DFG), Project number 511060241. Funders had no influence on study design, collection, analysis and interpretation of data.

## Supplementary Figures

**Figure S1.**
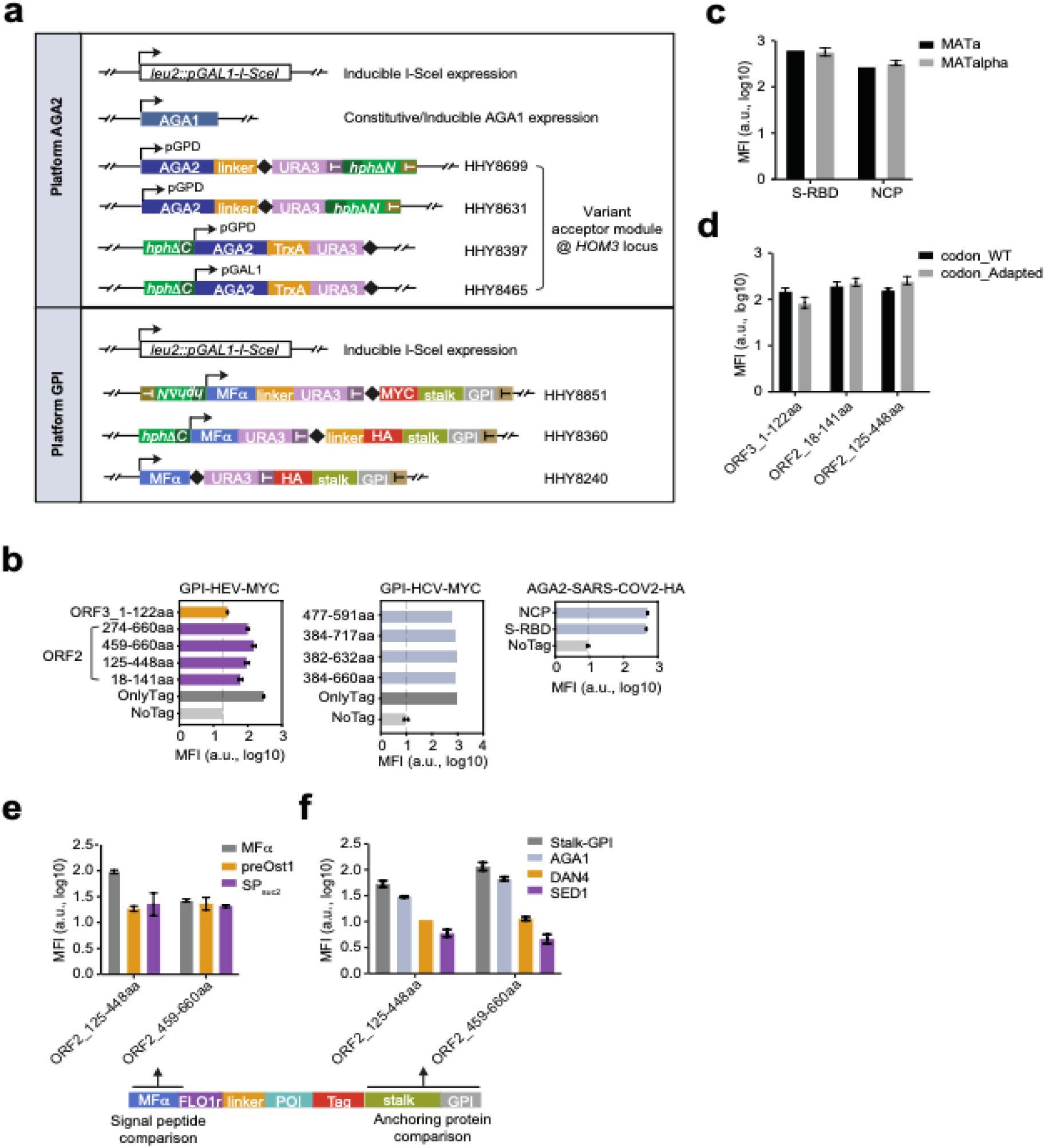
Construction and optimization of YSD strains expressing various viral antigens. (**a**) Schematic representation of the acceptor strains used for the different display platforms. The I-SceI motif is indicated by a black diamond. (**b**) Surface expression levels of various antigens displayed using the AGA2- and GPI-based platforms. Antigens were derived from hepatitis E virus (HEV), hepatitis C virus (HCV), and SARS-CoV-2. Strains expressing only the epitope tag served as positive controls, and strains lacking any tag served as negative controls. Median fluorescence intensity (MFI; log10-transformed) of the epitope tag is shown. (**c**) Effect of mating type on surface display efficiency. Median fluorescence intensity (MFI; log10-transformed) of the epitope tag is shown. (**d**) Effect of codon usage on antigen expression levels. Median fluorescence intensity (MFI; log10-transformed) of the epitope tag is shown. (**e**) Comparison of alternative signal peptides for surface display efficiency. Median fluorescence intensity (MFI; log10-transformed) of the epitope tag is shown. (**f**) Evaluation of different anchor proteins for optimal surface expression. Median fluorescence intensity (MFI; log10-transformed) of the epitope tag is shown.

**Figure S2.**
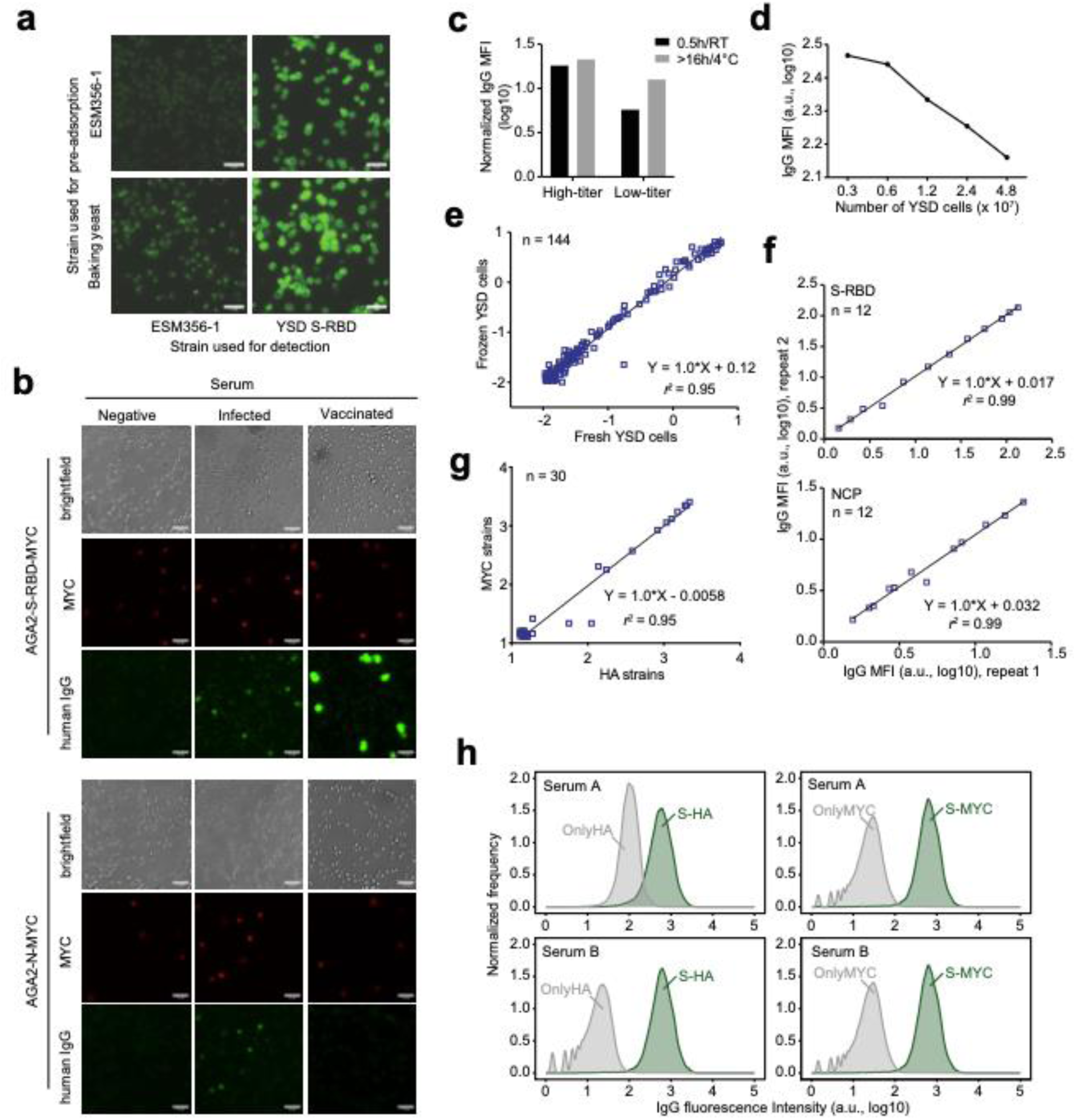
Validation and optimization of the YSD-FCM antibody detection workflow. (**a**) Effect of yeast pre-adsorption strain on serum reactivity detected by fluorescence microscopy. A SARS-CoV-2-positive serum was preadsorbed with either ESM356-1 or supermarket baking yeast, and residual reactivity was examined by fluorescence microscopy using ESM356-1 or YSD cells displaying S-RBD. Scale bar, 15 μm. (**b**) Detection of antibodies against S-RBD and NCP in human sera visualized by fluorescence microscopy. Images were acquired using identical exposure settings. Scale bar, 15 μm. (**c**) Effect of serum incubation time on detection sensitivity. Median fluorescence intensity (MFI; log10-transformed) of human IgG on the S-RBD YSD strain is normalized to that of the MYC-tag control strain. (**d**) Influence of YSD cell density during serum incubation on fluorescence intensity. Median fluorescence intensity (MFI; log10-transformed) of human IgG is shown. (**e**) Comparison of antibody detection using freshly cultured and frozen YSD cells. Median fluorescence intensity (MFI; log10-transformed) of human IgG on the S-RBD YSD strain is normalized to that of the MYC-tag control strain. (**f**) Reproducibility of the YSD-FCM assay. Correlation of two independent measurements (repeats 1 and 2) of anti-S-RBD and anti-NCP antibody levels in human sera (n = 12). Median fluorescence intensity (MFI; log10-transformed) of human IgG is shown. (**g**) Comparison of antibody detection performance between S-RBD-MYC and S-RBD-HA YSD strains. Median fluorescence intensity (MFI; log10-transformed) of human IgG is shown. (**h**) Recognition of the HA tag by non-specific antibodies present in human samples.

**Figure S3.**
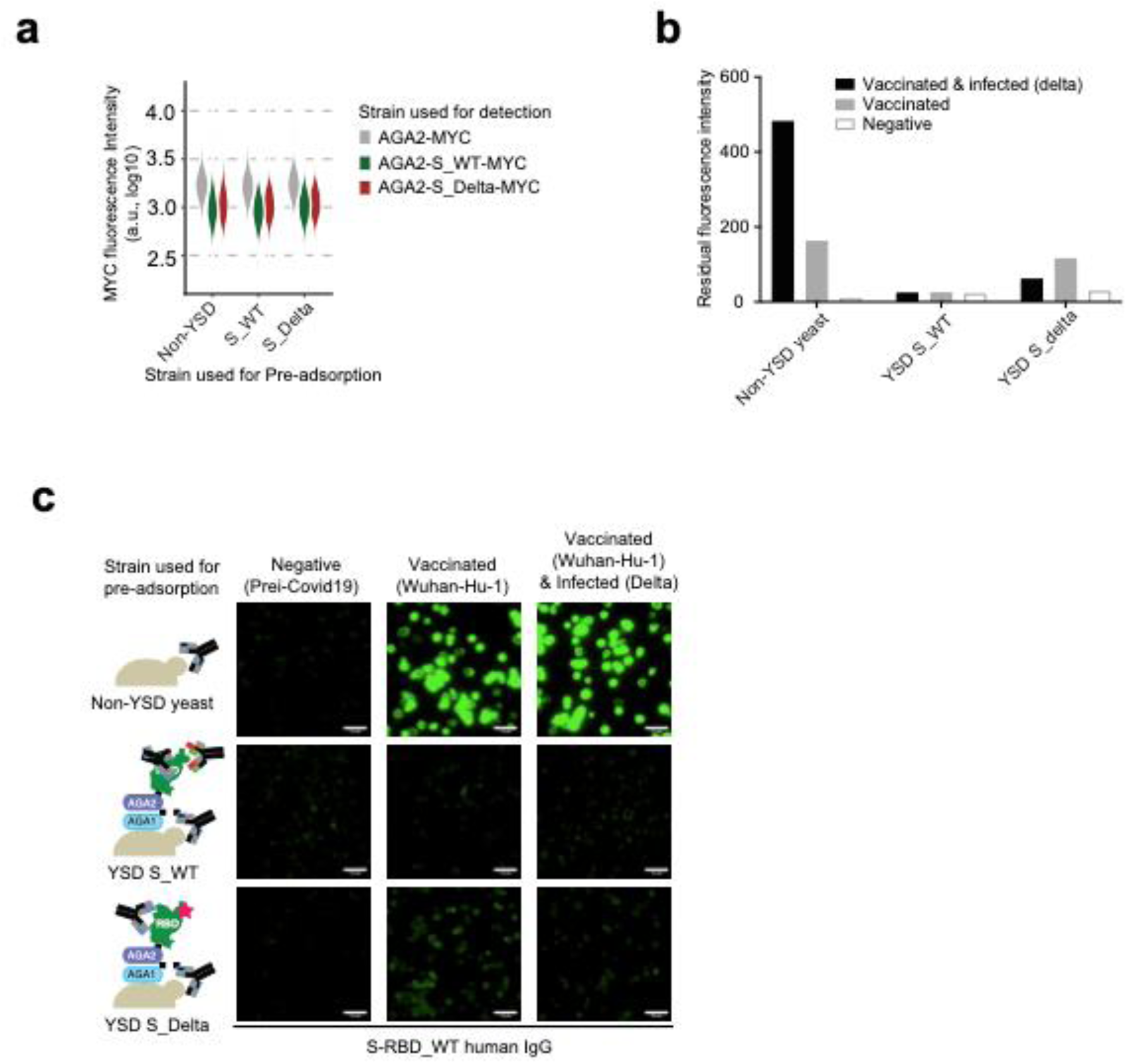
Variant-specific antibody depletion using YSD strains. (**a**) Violin plots showing fluorescence intensity of the MYC epitope tag on AGA2-MYC, AGA2-S_WT, and AGA2-S_Delta strains used for variant-specific antibody depletion experiments. (**b**) Residual fluorescence intensity after variant-specific antibody depletion, measured by flow cytometry. Median fluorescence intensity (MFI; log10-transformed) of human IgG is shown. (**c**) Variant-specific antibody depletion visualized by fluorescence microscopy using YSD strains. Images were acquired using identical exposure settings. Scale bar, 15 μm.

**Figure S4.**
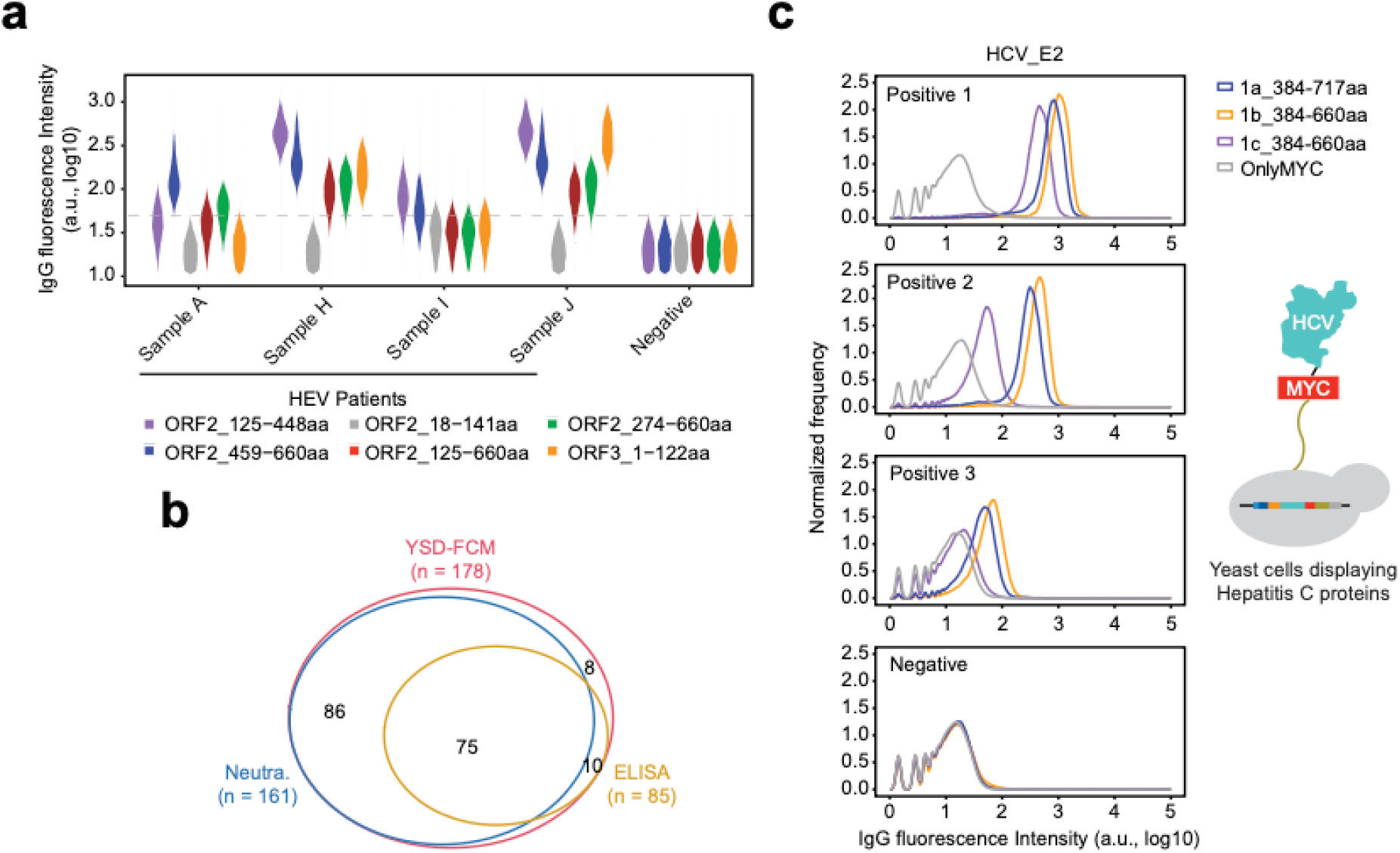
Quantitative detection of antibodies against hepatitis viruses using yeast surface display and flow cytometry (YSD-FCM). (**a**) Violin plot showing human IgG fluorescence intensity for different HEV-derived fragments measured by flow cytometry. YSD strains displaying individual fragments were incubated with serum samples from HEV-infected and HEV-negative individuals. (**b**) Venn diagram illustrating the overlap of serum samples analyzed by different assays: 178 serum samples were analyzed by YSD-FCM, 85 samples were analyzed by ELISA, and 161 samples were analyzed using a neutralization assay. (**c**) Detection of antibodies against hepatitis C virus (HCV) envelope glycoprotein E2 in serum samples from HCV-infected and HCV-negative individuals using flow cytometry.

**Figure S5.**
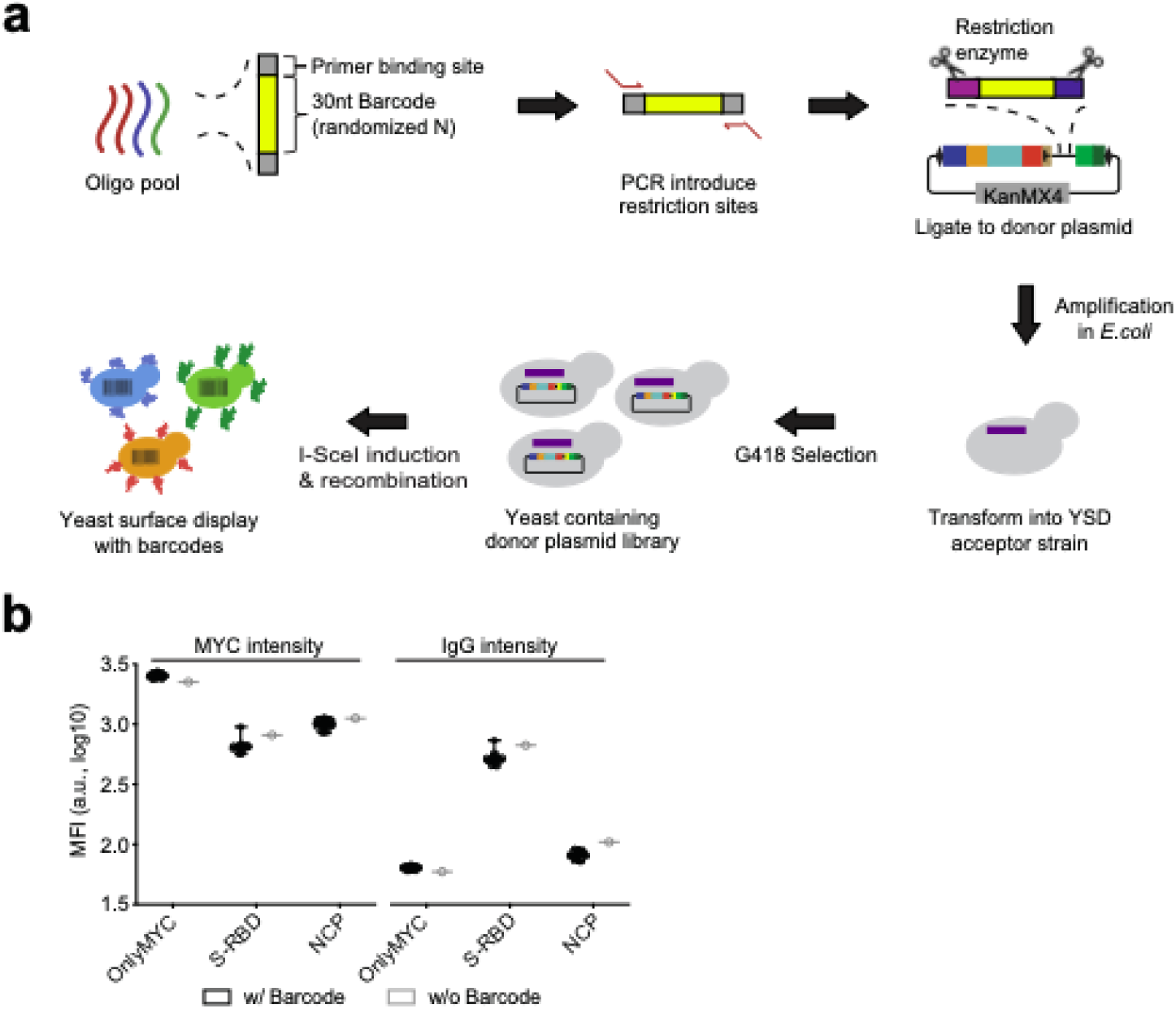
Barcoded YSD libraries used for SeroSeq. (a) Workflow for constructing barcoded YSD libraries. A pool of unique barcode oligonucleotides was cloned into donor plasmids carrying the respective POI (SARS-CoV-2 S-RBD, NCP, or MYC-tag).(b) Comparison of MYC-tag expression and IgG-binding levels between barcoded and non-barcoded strains.

**Figure S6.**
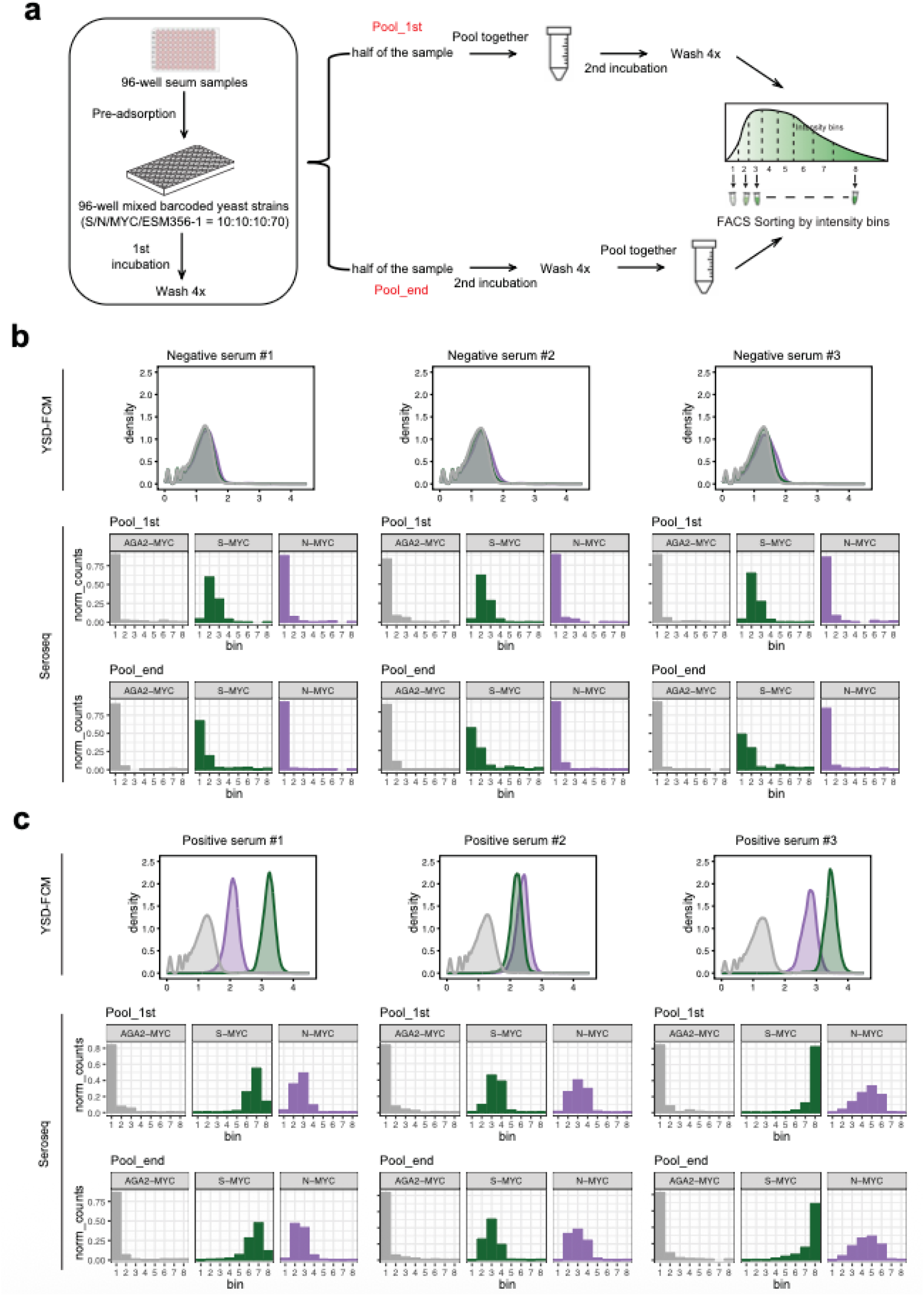
Comparison of Pool_1st and Pool_end pooling strategies in SeroSeq. (**a**) Workflow of Pool_1st and Pool_end pooling strategies. (**b**) Representative results obtained from three seronegative samples. (**c**) Representative results obtained from three seropositive samples.

**Figure S7.**
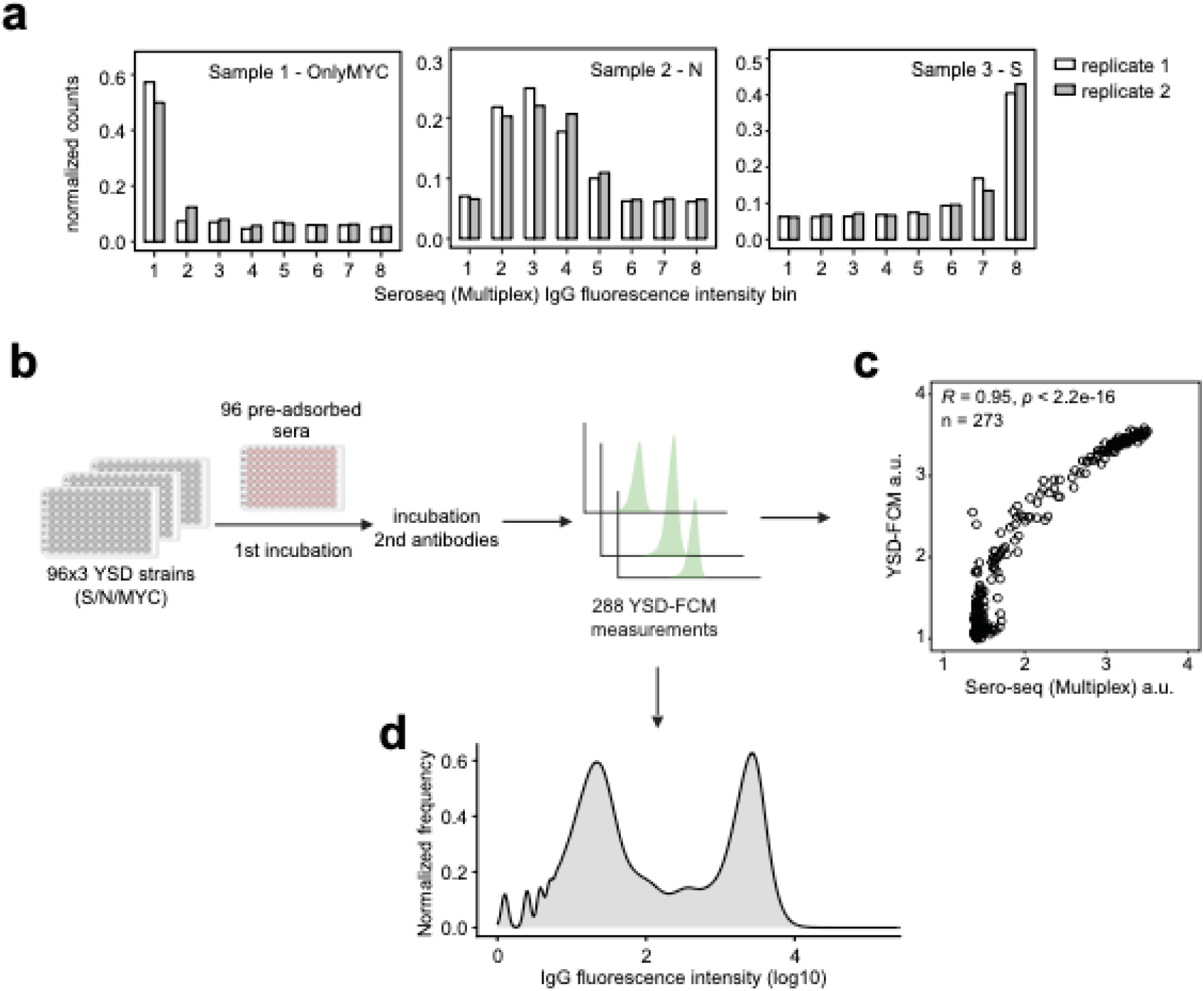
Validation of SeroSeq. (**a**) Duplicate analysis of the SeroSeq assay. Representative bar plots show antibody profiles from negative, infected, and vaccinated donors tested in duplicate using different barcodes. (**b**) Workflow for matched single-sample YSD-FCM assays. 96 serum samples were analyzed individually in 96-well format by YSD-FCM using SARS-CoV-2 S-RBD, NCP, or MYC-tag strains. Each plate contained one YSD strain across all wells, enabling matched measurement of antigen-specific antibody binding for every serum sample. This workflow generated 288 measurements across three antibodies. (**c**) Correlation between multiplexed SeroSeq and single-sample YSD-FCM measurements. Normalized read counts for each replicate are shown as bar plots. (**d**) Combined YSD-FCM dataset for comparison with multiplexed SeroSeq measurements. Single-sample measurements obtained from individual YSD-FCM assays across all 96 serum samples were combined into one dataset.

